# Evolutionary Dynamics of *Paramecium* Mitochondrial Genomes

**DOI:** 10.64898/2026.05.26.727735

**Authors:** Ravina Telkar, Farhan Ali, Samuel F. Miller, Jiahao Ni, Lydia Bright, John P. DeLong, Kristi L. Montooth, Sascha Krenek, Masahiro Fujishima, Kenji Nanba, Michael Lynch

## Abstract

Ciliates are a diverse group of single-celled eukaryotes that can exhibit a wide range of genetic diversity within morphologically indistinguishable species. However, they are still not well studied as their mechanisms of speciation and the extent of diversification remain unknown. Mitochondrial genomes offer an effective framework for resolving species relationships and evolutionary changes. Here, we analyzed a globally sampled dataset of *Paramecium* to understand the evolution of mitochondrial genomes in ciliates. Phylogenetic analysis of linear mitochondrial genomes shows the presence of cryptic diversity beyond the *P. aurelia* complex, with *P. bursaria* lineage appearing as a deeply diverging out-group. Protein-coding genes are largely conserved, with limited rearrangements, and some ciliate-specific genes appear to be missing in *P. bursaria*. Population genetic analysis show little to no evidence of recombination along with substantial differences in effective population size across species. Patterns of molecular evolution also indicate purifying selection as the predominant force, the strength of which is at least as strong as in the nucleus and consistent with mitochondrial effective population sizes that are similar or larger than those of the nucleus. Across the functional groups, the electron transport chain and ribosomal genes are highly constrained, while ciliate-specific *ymf* genes show reduced efficacy of selection compared to the others. These findings offer a basis for connecting mitochondrial variation to evolutionary divergence, functional constraint, and speciation in microbial eukaryotes.

## Introduction

Mitochondrial genomes evolve under a population-genetic environment that is fundamentally distinct from that of the nuclear genome. For instance, reduced efficacy of purifying selection in mitochondrial genes of animals, relative to nuclear genes, can be attributed to uniparental inheritance and the absence of recombination (Neiman & Taylor, 2009; Havird & Sloan, 2016). Genome-wide linkage effects arise as a result of low or no recombination, decreased effective population sizes, and are further impacted by higher rates of mutation (Lynch & Blanchard, 1998; Lynch et al., 2006). These characteristics are predicted to decrease the effectiveness of selection in eliminating deleterious mitochondrial mutations and promoting fixations of the beneficial ones, even in species with large nuclear populations. On the other hand, we know that mitochondrial genome recombination has been reported in plants (e.g. *Arabidopsis thaliana*) and yeast (e.g. *Saccharomyces cerevisiae*), indicating that evolutionary dynamics of the mitochondrial genome can vary widely among different phylogenetic lineages (Piganeau et al., 2004; Barr et al., 2005; Havird & Sloan, 2016; Fritsch et al., 2014). These variations demonstrate the necessity to examine how variations in recombination and modes of inheritance affect the evolution of the mitochondrial genome in different eukaryotic lineages.

Ciliates such as *Paramecium* are powerful model systems for studying population genetics and genome evolution, owing to the extensive intraspecific diversity, global distribution, and their well-characterized genetics (Sonneborn, 1975; Finlay, 2002; Fenchel & Finlay, 2004; Johri et al., 2017). Studies on cryptic lineages and species complexes help us understand speciation and divergence in microbial eukaryotes (Sonneborn, 1975; Barth et al., 2008; Tarcz et al., 2012; Melekhin et al., 2022). However, key aspects of how evolutionary dynamics shape genome variation and species formation in protists remain comparatively underexplored relative to multicellular systems (Seehausen et al., 2014) and well-studied unicellular model systems such as yeast (e.g. *Saccharomyces cerevisiae*) (Liti et al., 2009). Studying mitochondrial population genomics in ciliates enables us to understand whether mitochondrial biology contributes to cryptic speciation and how evolutionary forces shape genome evolution across lineages, while facilitating comparisons of genome organization among closely related species due to the ease of sequencing and alignment of mitochondrial genomes.

In *Paramecium*, the mitochondrial genomes are streamlined and contain genes essential for oxidative phosphorylation and mitochondrial translation. Patterns of molecular evolution in mitochondrial genes are closely linked to those of their nuclear-encoded interacting partners and are often expected to reflect strong purifying selection despite constraints imposed by their population-genetic structure, including reduced effective population size, limited recombination, and strong genome-wide linkage (Lynch et al., 2006; Havird & Sloan, 2016). Mitochondrial genomes can harbor lineage-specific genes, such as the ciliate-specific *ymf* family, which evolve rapidly, but the relative contributions of relaxed constraint and positive selection remain unresolved (Johri et al., 2019). Mitochondrial genome evolution cannot be easily understood from a single-species perspective because important population-genetic parameters differ between eukaryotic lineages. Instead, comparative population-genomic analyses are necessary to identify the roles of mutation, linkage, and selection (Barr et al., 2005; Smith & Keeling, 2015).

The evolutionary dynamics of mitochondrial genomes in *Paramecium* have been characterized by earlier research, identifying strong purifying selection, conserved gene order, and no detectable recombination across a limited set of species (Johri et al., 2019). However, restricted sampling limits the generality of these findings and leaves variation across lineages unresolved. In order to provide further population-level resolution and phylogenetic depth, we expand this framework by analyzing 315 mitochondrial genomes from 22 globally distributed species, including cryptic lineages both inside and outside the *P.aurelia* complex. This expanded sampling enables robust inference of selection, including tests for positive selection and relaxed constraint, allowing us to distinguish the evolutionary forces acting on genes such as *ymf*. It also allows us to examine differences in evolutionary rates among lineages, compare functional patterns with nuclear-encoded genes, and characterize genome structure. In addition, our dataset helps identify cryptic and potentially new species. The dynamics of mitochondrial evolution across lineages are better understood from these analyses.

## Results

### Mitochondrial phylogeny of globally sampled *Paramecium* reveals cryptic lineages

The Ciliata frequently hide significant genetic diversity that cannot be identified by morphology alone. One classic example is *Paramecium aurelia*, which was shown through genetic studies to comprise multiple reproductively isolated sibling species (Sonneborn, 1975). This pattern was later confirmed by genomic analyses across the genus (Johri et al., 2017; McGrath et al., 2014) and observed in other ciliates like *Tetrahymena* (Zhou et al., 2022). To discover cryptic diversity and lineage relationships in *Paramecium*, we assembled mitochondrial genomes from a globally distributed collection of isolates from Africa, Asia, North and South America, Europe, and the Pacific region (Fig. 1A; Supp. Table S1). We obtained high-confidence mitochondrial assemblies for 315 samples representing 22 species, including the 11 species examined in previous mitochondrial phylogenies. Members of the *P. aurelia* complex exhibited very consistent genome sizes (∼39–40 kb), whereas more distantly related taxa like *P. bursaria* had larger mitochondrial genomes (∼44 kb). The range of mitochondrial genome sizes was ∼30–48 kb.

**Fig. 1.**
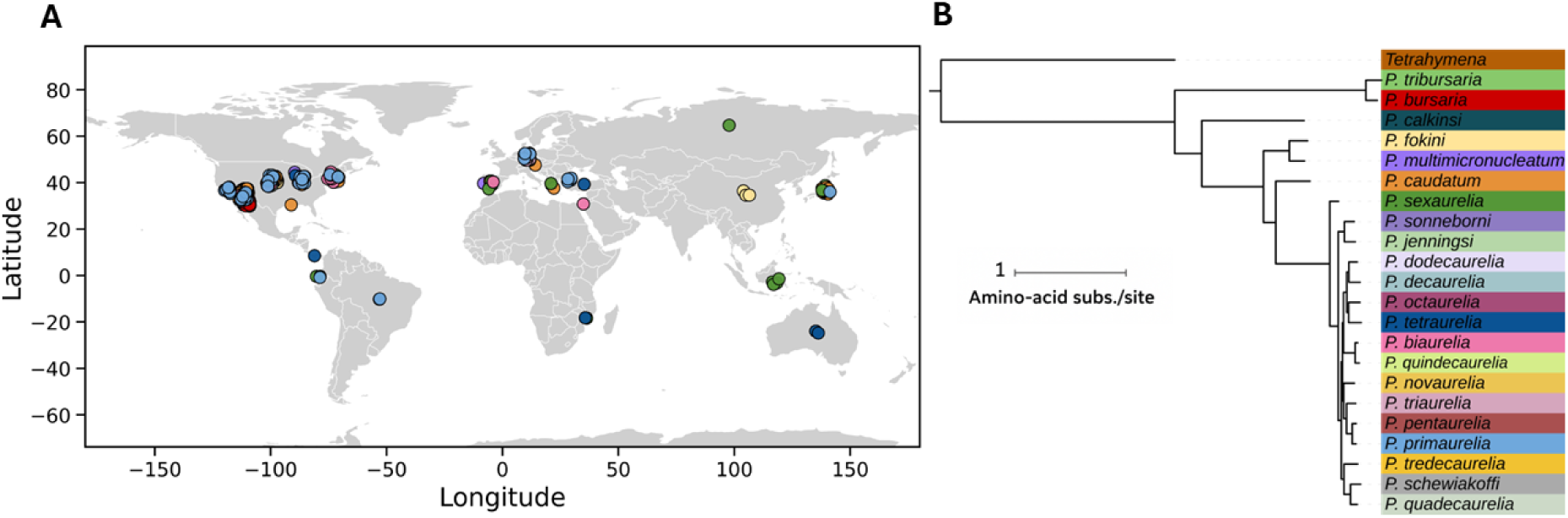
Global sampling and evolutionary relationships of *Paramecium* species. (A) Global distribution of *Paramecium* species analyzed in this study. (B) Phylogenetic relationships among *Paramecium* species based on mitochondrial gene analysis.

A well-resolved phylogeny for 22 *Paramecium* species with *Tetrahymena* as an outgroup was obtained from the concatenated mitochondrial alignment of 42 genes, including ribosomal, electron-transport, and ciliate-specific *ymf* genes (Fig. 1B), inferred using RAxML-NG under the LG+G substitution model with 100 bootstrap replicates (Kozlov et al., 2019). The *P. bursaria* lineage formed a deeply branching outgroup, while *P. multimicronucleatum* and *P. fokini* form a well-supported sister pair with *P. caudatum* outside the *P. aurelia* complex. *P. sexaurelia* is strongly placed as sister to the remaining *P. aurelia* species, consistent with previous phylogenetic analyses (Johri et al., 2019). We confirmed the concatenated topology using independent gene-tree analyses, with the majority of genes supporting the overall relationships (Suppl. Table S2), although support for specific relationships varies across genes. Our expanded sampling recovers patterns consistent with previously reported cryptic lineages in *Paramecium*, where morphologically defined species can form deeply diverged mitochondrial lineages likely representing distinct species (Johri et al., 2019). Notably, the previously reported isolate *P. multimicronucleatum*-M04 clusters with *P. fokini*, while C026 remains distinct from other *P. caudatum* samples (Johri et al., 2019). We also recover a close relationship between *P. biaurelia* and *P. quindecaurelia* and confirm cryptic diversity within the *P. bursaria* and *P. multimicronucleatum* complexes (Hori et al., 2005; Melekhin et al., 2022). Together, these results support and refine existing evidence for cryptic speciation in *Paramecium*.

### Architecture and evolutionary divergence of *Paramecium* mitochondrial genomes

Mitochondrial genomes of ciliates display considerable structural diversity, with substantial variation in genome size, gene order, and gene content reported in species such as *Tetrahymena thermophila* and *Euplotes* (*E. minuta* and *E. crassus*) (Brunk et al., 2003; de Graaf et al., 2009). To determine how mitochondrial genome architecture is conserved in *Paramecium* and where lineage-specific divergence has taken place, we looked at patterns of gene presence, genome organization, gene length variation, and nucleotide composition across *Paramecium* species.

### Mitochondrial gene content and genome organization

Mitochondrial gene-content analysis identified nearly all genes previously reported in *Paramecium* mitochondrial genomes, including 15 oxidative phosphorylation genes, 10 ribosomal protein genes, and multiple *ymf* and other uncharacterized ORFs (Fig. 2A), consistent with previous annotations (Arnaiz et al., 2019). Overall, the gene content is well conserved. Except for *rpS13*, which is lacking in *P. tredecaurelia*, 45 protein-coding genes are shared by *P. aurelia* species. Variation is primarily restricted to rapidly evolving *ymf* genes in more divergent lineages, such as *P. bursaria* and *P. fokini*. Nevertheless, some apparent absences in *P. fokini* may suggest an incomplete mitochondrial assembly (∼30 kb versus ∼40 kb in other species). The constant retention of *ymf* genes throughout the genus suggests that they are important but very divergent components of ciliate mitochondrial genomes, despite their unknown functions. This is supported by apparent gaps in *P. bursaria*, that might correspond to highly divergent, ciliate-specific *ymf* genes that are difficult to identify based only on sequence similarity, indicating that these areas have concealed homology rather than actual gene loss (Supp. Note 1).

**Fig. 2.**
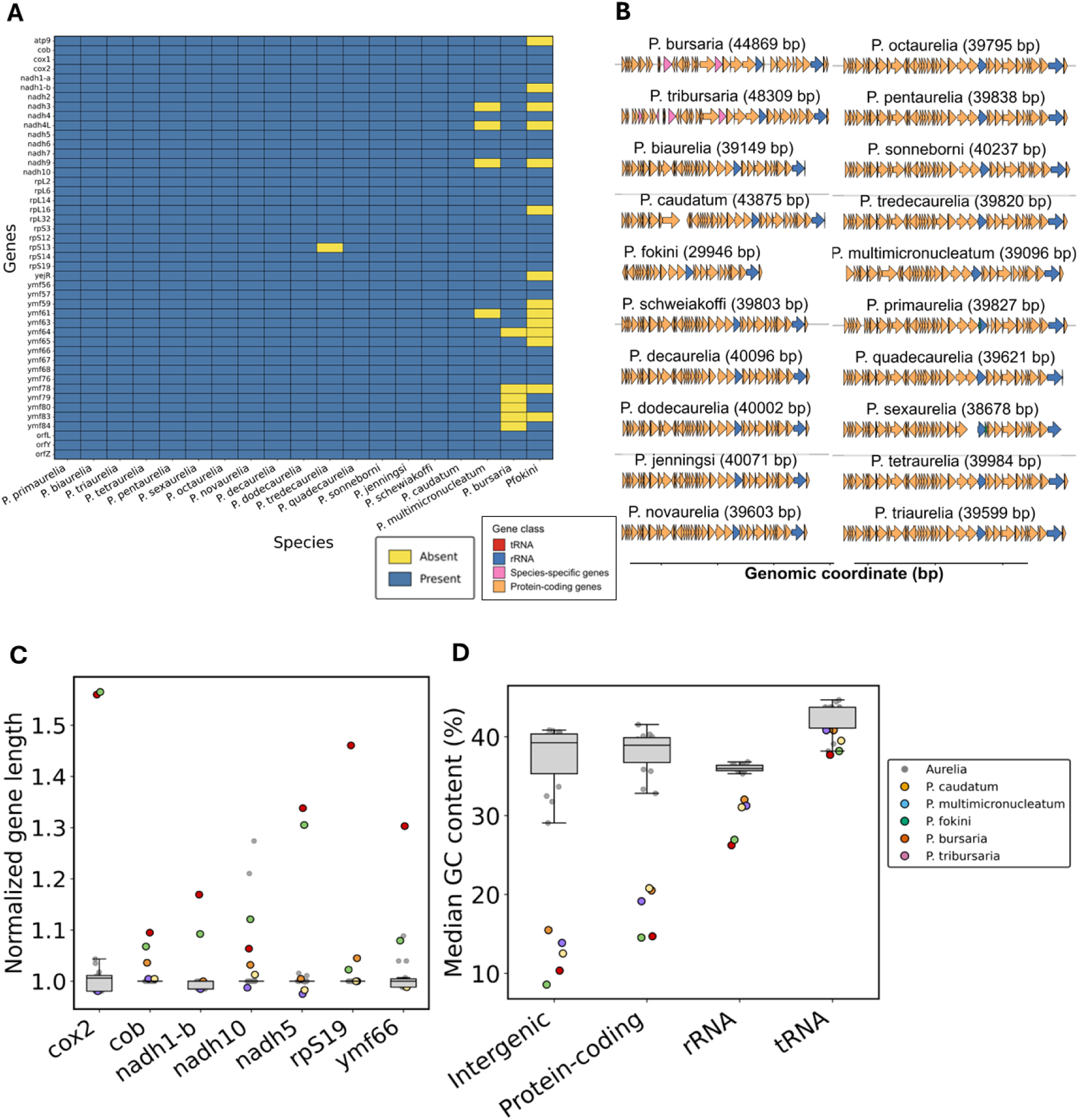
Conservation and lineage-specific divergence of *Paramecium* mitochondrial genomes. (A) Presence–absence matrix of mitochondrial genes across *Paramecium* species. (B) Mitochondrial genome organization showing conserved gene order with limited rearrangements. (C) Normalized gene length variation, with greater variability in outgroup species. (D) GC content varia-tion across genomic features, with elevated GC in *P. aurelia* and consistently high GC in rRNA and tRNA genes.

Consistent with gene content, mitochondrial genome organization is also broadly conserved (Fig. 2B), with most genes maintaining the same relative order across species. Key RNA genes, including *tRNA-Tyr*, *tRNA-Phe*, *tRNA-Trp*, and SSU and LSU rRNA, also occur in similar genomic positions, whereas limited rearrangements are observed primarily in the *bursaria* lineage, where genes such as *tRNA-Trp*, SSU rRNA, *rpL14*, and *rpL16* occupy different positions. This pattern is consistent with the greater divergence of *P. bursaria* relative to the *P. aurelia* complex (Serra et al., 2021). The conservation of gene order across the genus suggests that they are subject to strong constraints, possibly related to RNA processing and coordinated transcription.

### Mitochondrial gene length and GC content variation

Although mitochondrial gene content and order are generally conserved, we observed a range of mitochondrial genome sizes across species; the longest genome is nearly 1.5 times the length of the shortest genome. These variations in genome sizes could result from variations in gene length that maintain essential mitochondrial functions. We analyzed mitochondrial gene lengths across species to test whether *Paramecium* shows similar trends. Within the *P. aurelia* complex, gene lengths are highly conserved. Most genes cluster closely around their median size (Fig. 2C; Supp Fig. S1A). Unlike species within this complex, species outside it, particularly those in the *P. bursaria* lineage, show marked expansions in a subset of genes, including *nadh1-b*, *nadh10*, *nadh5*, *rpS19*, and *ymf66*, reaching approximately 1.5–4× the genus-wide median. Together, these findings strongly suggest gene-specific divergence, rather than genome-wide changes in mitochondrial gene length, across the genus.

Variation in mitochondrial GC content provides insight into evolutionary constraints and mutation biases. Mitochondrial nucleotide composition varies widely across eukaryotes, and many ciliates have strongly AT-rich mitochondrial genomes, including *Tetrahymena* (18–22% GC) and *Euplotes* (Brunk et al., 2003; de Graaf et al., 2009; Johri et al., 2019). In *Paramecium*, GC content shows a clear lineage-specific pattern (Supp. Table S3; Fig. 2D). In accordance with AT-rich mitochondrial genomes, distantly related species such as *P. bursaria*, *P. tribursaria*, *P. caudatum*, *P. multimicronucleatum*, *P. fokini*, and *P. calkinsi* have lower GC content (14–20%). The *P. aurelia* complex, on the other hand, show elevated GC levels (32–42%). Both intergenic areas and protein-coding genes exhibit this difference (Supp Fig. S1B), with *P. aurelia* species often exceeding 39% GC, while outgroup species remain ≤22%. These concordant patterns indicate a substantial genome-wide compositional shift in the *P. aurelia* lineage.

Structural RNA genes show a distinct pattern. rRNA and tRNA genes remain relatively GC-rich even in AT-rich genomes. tRNA GC content is consistently high (∼40–46%) across species, whereas rRNA GC content ranges from ∼26–32% in non-*aurelia* species to ∼36–37% in *aurelia* species. Thus, the genome-wide GC pattern observed in protein-coding genes, that might be driven by mutation bias, is not fully reflected in mitochondrial RNAs. One possible explanation is that structural requirements of rRNAs and tRNAs favor higher GC content to maintain stable RNA secondary structures, since GC base pairing contributes to RNA stability (Sueoka, 1993; Goodenbour & Pan, 2006; Gray, 2003). Similar GC enrichment in tRNAs and rRNAs is observed across diverse mitochondrial genomes, including *Homo sapiens* (Anderson et al., 1981), *Drosophila melanogaster* (Clary & Wolstenholme, 1985), and *Saccharomyces cerevisiae* (Foury et al., 1998).

### Patterns of Nucleotide Diversity Across Mitochondrial Site Classes

Levels of sequence variation differ across functional site classes because mutations at some positions have stronger fitness consequences. Sites under stronger functional constraint are therefore expected to show lower variation. To characterize mitochondrial variation in *Paramecium*, we estimated nucleotide diversity (*π*) for each species, as the average number of nucleotide differences per site between sequences (Nei, 1987; Lynch, 2007), averaged over all sample pairs. Nucleotide diversity varies significantly among site classes, consistent with differences in functional constraint (Table 1).

**Table 1.** Nucleotide diversity across mitochondrial site classes in *Paramecium*. Species are ordered by overall nucleotide diversity (*π*). Values marked as “–” indicate missing data due to incomplete genome assembly.

| Species | Overall | tRNA | rRNA | 0-fold | Intergenic | 2-fold | 4-fold |
| --- | --- | --- | --- | --- | --- | --- | --- |
| *P. caudatum* | 0.0665 | 0.0018 | 0.0236 | 0.0334 | 0.1099 | 0.1057 | 0.1234 |
| *P. sexaurelia* | 0.0546 | 0.0000 | 0.0071 | 0.0111 | 0.0811 | 0.1356 | 0.1574 |
| *P. fokini* | 0.0249 | 0.0141 | 0.0100 | 0.0193 | 0.0302 | 0.0390 | 0.0581 |
| *P. primaurelia* | 0.0143 | 0.0024 | 0.0095 | 0.0027 | 0.0116 | 0.0385 | 0.0401 |
| *P. bursaria* | 0.0105 | 0.0000 | 0.0020 | 0.0052 | 0.0135 | 0.0165 | 0.0252 |
| *P. quindecimurelia* | 0.0104 | — | — | 0.0052 | 0.0063 | 0.0107 | 0.0073 |
| *P. biaurelia* | 0.0075 | 0.0019 | 0.0001 | 0.0179 | 0.0049 | 0.0266 | 0.0271 |
| *P. tetraurelia* | 0.0065 | 0.0000 | 0.0005 | 0.0027 | 0.0080 | 0.0140 | 0.0134 |
| *P. triaurelia* | 0.0059 | 0.0066 | 0.0010 | 0.0021 | 0.0102 | 0.0173 | 0.0216 |
| *P. tribursaria* | 0.0050 | 0.0000 | 0.0002 | 0.0033 | 0.0074 | 0.0091 | 0.0093 |
| *P. multimicronucleatum* | 0.0040 | 0.0000 | 0.0006 | 0.0027 | 0.0042 | 0.0099 | 0.0055 |
| *P. quadecaurelia* | 0.0005 | 0.0000 | 0.0002 | 0.0007 | 0.0009 | 0.0013 | 0.0019 |

The rRNA and tRNA regions of structural RNA genes have extremely little variation, and these genes have the lowest nucleotide diversity. Their crucial functions in mitochondrial translation, where ribosome structure and precise codon recognition depend on sequence integrity, are reflected in this strong conservation (Gray et al., 1999; Shajani et al., 2011; Suzuki & Suzuki, 2014). Purifying selection against mutations that modify amino acids is reflected in the constraint of nonsynonymous (0-fold degenerate) sites in protein-coding genes in comparison to synonymous sites. Diversity increases at partially degenerate (2-fold) sites and is highest at 4-fold degenerate sites, which may be largely neutral. Intergenic regions show intermediate levels of variation. Overall, nucleotide diversity follows the hierarchy: tRNA ≈ rRNA *<* 0-fold *<* intergenic *<* 2-fold *<* 4-fold

Earlier work on ciliate mitochondria shows similar trends. Highest nucleotide diversity at four-fold degenerate sites (*π*_4_), used as a measure of neutral change, appears in *P. sexaurelia* and *P. caudatum*. Our findings, 0.157 and 0.123, agree with prior results 0.169 and 0.127 (Johri et al., 2017). Such levels are consistent with high mitochondrial polymorphism in ciliates. Nucleotide variation in *Paramecium* matches or exceeds levels seen in many microbial eukaryotes; for example, in *Saccharomyces cerevisiae*, mitochondrial diversity usually measures *π* = 0.001–0.003 (Liti et al., 2009; Peter et al., 2018). Some limitations should be noted. For certain species (e.g., *P. multimicronucleatum*, *P. quadecaurelia*, and *P. biaurelia*), only one complete mitochondrial genome is available, with additional samples incomplete, reducing informative sites. The mitochondrial genome of *P. quindecaurelia* is incomplete due to insufficient sequence coverage and/or incomplete assembly, resulting in missing regions (such as rRNA and tRNA genes) and therefore missing estimates for some site classes. Even so, trends in genetic variation stay largely unchanged among species.

### Limited Evidence for Recombination in *Paramecium* Mitochondria

Recombination plays a central role in genome evolution by breaking linkage disequilibrium among sites and thereby enhancing the efficacy of natural selection. In its absence, genomes are expected to evolve as effectively nonrecombining units, with strong linkage enhancing the effects of background selection and genetic drift. Recombination in mitochondrial genomes can happen through processes such as transient heteroplasmy, biparental inheritance, or mitochondrial fusion, despite the common assumption that mitochondrial genomes are clonal. Ciliate mitogenomes show contrasting organization, with extensive rearrangements in *Colpoda* despite no detectable recombination (Zhang et al., 2024). We observed that gene order is highly conserved in *Paramecium*. These contrasting observations suggest that genome architecture alone does not provide direct evidence for or against recombination; this motivated a direct test for recombination in *Paramecium*.

To evaluate recombination, we applied the PHI (Pairwise Homoplasy Index) test, which detects excess homoplasy among polymorphic sites (Bruen et al., 2006) (Fig. 3A). Most gene–species combinations showed no significant signal or insufficient polymorphism, indicating limited power to detect recombination. Occasional signals were rare and not reproducible across genes. Additional tests, including the Neighbor Similarity Score (NSS) and Maximum Chi-Square (MaxChi^2^) test (Bruen et al., 2006), yielded consistent results (Supp. Fig. S2). Because mitochondrial recombination, if it occurs, is anticipated to occur mostly as short-tract, nonreciprocal events that may escape detection by conventional recombination tests, we also evaluated the potential of localized, nonreciprocal exchange using LDhat under a gene-conversion framework (McVean et al., 2002). Recombination-rate estimations were close to zero across species, and LD-distance correlations and permutation-based statistics were not significant (Fig. 3B, Supp. Table S4). Because recombination cannot be estimated independently of effective population size, we report the population-scaled recombination parameter (4*N_e_r*), where *r* is the per-site recombination rate, to enable comparisons across species. Estimates of 4*N_e_r* were near zero, and occasional non-zero values were not supported by other LD-based statistics, indicating that these likely reflect estimation noise rather than true recombination. Only *P. biaurelia* showed nominally significant composite-likelihood values, although these were not supported by other statistics. Weak and inconsistent signals were also observed in *P. sexaurelia* and *P. primaurelia*, but these were not reproducible across methods and do not provide robust evidence for recombination. All other species showed no evidence of gene conversion. This conclusion supports a model of effectively clonal mitochondrial inheritance and is consistent with classical genetic studies indicating strict uniparental inheritance and a lack of mitochondrial fusion (Adoutte et al., 1979).

**Fig. 3.**
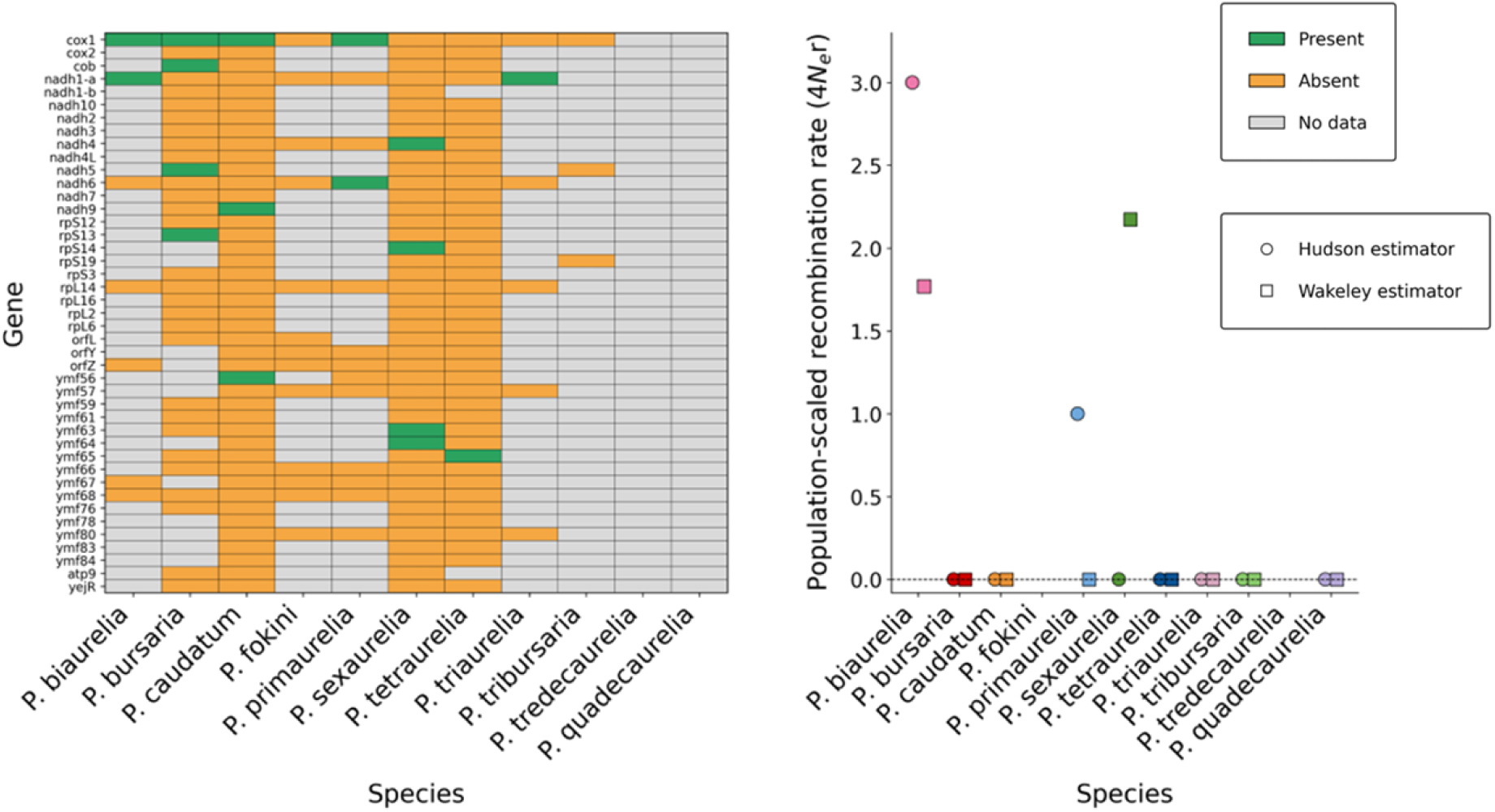
Evidence for mitochondrial recombination across *Paramecium* species. (A) PHI (Pairwise Homoplasy Index) permutation test across mitochondrial genes and species. Green indicates nominal significance (*p <* 0.05), orange indicates non-significance (p *≥* 0.05), and gray denotes limited polymorphism data. (B) LDhat gene-conversion estimates of the population-scaled recombination rate (4*Ne*r) at the smallest *θ* value for each species. Both Hudson (circles) and Wakeley (squares) estimators yield rates that are zero or near zero across taxa.

### Effective population size shapes the efficacy of purifying selection across *Paramecium* species

Mitochondrial genomes in *Paramecium* provide a valuable paradigm for investigating the relationship between genome-wide patterns of genetic evolution and variation in effective population size (*N_e_*). We have seen that silent sites have the highest nucleotide diversity in all species, which presumably reflects weakest functional constraints among all classes of sites. Furthermore, several aspects of mitochondrial genome architecture appears to be conserved, suggesting similar selection pressures across species. Mitochondrial genetic divergence does not increase with geographic distance, consistent with widespread dispersal in free-living microbial eukaryotes, indicating no isolation by distance and no geographic structuring of neutral mitochondrial variation (Finlay, 2002; Fenchel & Finlay, 2004; Foissner, 2006). Assuming that mitochondrial mutation rate is also largely invariant among these closely related species, variation in synonymous diversity (*π_S_*) across species should reflect differences in long-term *N_e_*, in accordance with the neutral expectation *π_S_*= 2*N_e_µ*. Mean synonymous diversity (*π_S_*), calculated across genes for each species, varied widely among species (Fig. 4A; Supp. Table S5), with *P. sexaurelia* (0.237 ± 0.012) and *P. caudatum* (0.184 ± 0.009) exhibiting the highest values, indicative of larger effective population sizes, whereas species such as *P. biaurelia* (0.012 ± 0.003) show markedly lower diversity, consistent with smaller *N_e_*.

**Fig. 4.**
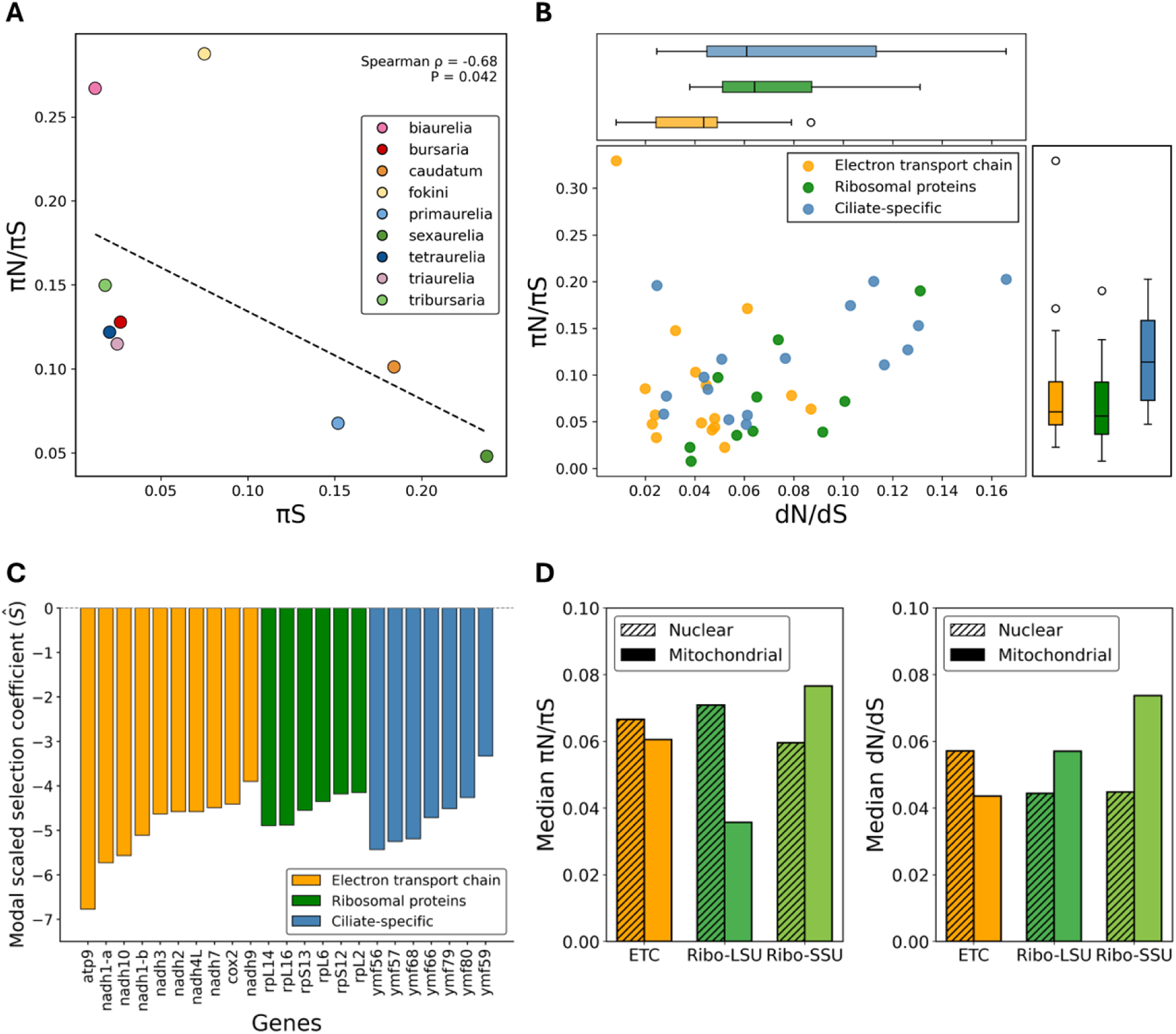
Relative effective population size and selection efficacy in *Paramecium*. (A) Synonymous diversity (*π_S_*), used as a proxy for long-term effective population size, and the efficacy of purifying selection (*π_N_ /π_S_*), showing stronger selection in species with higher inferred *N_e_*. (B) *π_N_ /π_S_* for within-species polymorphism and *d_N_ /d_S_* for between-species divergence were used to evaluate gene-level selection patterns across mitochondrial functional groups. The ratio of nonsynonymous to synonymous diversity for each gene was computed as *π_N_ /π_S_*, which was then summed across genes within each functional category using the median.The M8 model was used to estimate divergence-based selection, which permits *ω* (*d_N_ /d_S_*) to differ between sites within a same phylogeny. (C) Gene-wise modal scaled selection coefficients (*S*^^^; *S* = 2*Nes*). (D) Nuclear–mitochondrial comparison of selection efficacy across functional categories.

Species with large *N_e_*should exhibit a stronger efficacy of purifying selection. Consistent with this, we observed a strong negative correlation between the average of *π_N_ /π_S_* over genes and *π_S_* (Spearman *ρ* = −0.68, *P* = 0.042), after removing species with limited gene representation due to low sampling and incomplete genomes (such as *P. multimicronucleatum* and *P. quindecaurelia*). This was not merely an intrinsic correlation between a ratio and its denominator because we measured *π_N_ /π_S_* and *π_S_* on different sets of isolates. Together, these results reveal the differences in mitochondrial effective population size and consequently, in the efficacy of purifying selection on mitochondrial genomes across species of *Paramecium*.

### Selective constraint across mitochondrial genes and functional groups

Functional classes of mitochondrial genes show clear differences in selective constraint, as measured by *π_N_ /π_S_* (Fig. 4B). Ribosomal proteins (RP) and electron-transport chain (ETC) genes have the lowest values (medians = 0.0560 and 0.0605), consistent with strong purifying selection. Such conservation reflects their critical roles in energy production and protein synthesis inside mitochondria. Values remain well under 1; yet for lineage-specific *ymf* genes, *π_N_ /π_S_* reaches a median of 0.114, higher than in RP and ETC genes, hinting at weaker evolutionary constraint.

We estimated relative substitution rates at nonsynonymous and synonymous sites (*d_N_ /d_S_* = *ω*) using a maximum-likelihood framework (PAML Model M8; Supp Fig. S3), which allows *ω* to vary among sites while assuming a single distribution across the phylogeny. Under Model M8, *ω* represents the overall selective regime operating on each gene and represents a weighted average across site classes; values of *ω <* 1 indicate purifying selection, *ω* = 1 neutrality, and *ω >* 1 positive selection. *ω* remained below 1 for every gene, indicating extensive purifying selection. These constraints varied depending on the gene class: ETC genes had the lowest *ω* (median = 0.0436), followed by RP (0.0642) and *ymf* genes (0.0610) (Fig. 4B).

To further quantify the strength of selection, we converted the estimated probability distribution of *ω* from multi-species alignments to that of the scaled selection coefficients (*S* = 2*N_e_s*) based on the relation: 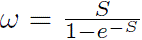 (Halpern and Bruno, 1998; Yang and Nielsen, 2008), and found that the mode of *S* representing the most common selection strength across sites inferred from all species together, was consistently negative across genes analyzed (Fig. 4C), indicating widespread purifying selection. This analysis was restricted to genes with nearly all sites in the beta-distributed class under the M8 model (*p*_0_ *>* 0.999), focusing on predominantly purifying selection. Strongest constraint was observed in core oxidative phosphorylation and ribosomal genes (e.g., *atp9*, *nadh1-a*, *nadh10*, *rpL14*, *rpL16*), whereas *ymf* genes showed less negative modal values (e.g., *ymf59*, *S* ≈ −3.3), consistent with weaker and more heterogeneous constraint. Although *ymf* genes remain predominantly under purifying selection, their elevated *π_N_ /π_S_*, *ω*, and less negative *S* indicate reduced constraint relative to other functional classes. However, gene-wide estimates can mask site-level heterogeneity, and average *ω <* 1 does not exclude positive selection at a subset of sites. Likewise, elevated *π_N_ /π_S_* may reflect relaxed constraint or mixed selective regimes. To distinguish between these possibilities, we computed neutrality indices (NI) and applied likelihood-based models that allow *ω* to vary among sites and across lineages, enabling detection of episodic positive selection despite overall gene-wide constraint.

### Limited evidence for positive selection across mitochondrial genes

To distinguish relaxed purifying selection from positive selection on individual genes, we computed neutrality indices (NI), defined as 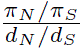. We estimated Branch-specific *d_N_* and *d_S_*using ancestral sequence reconstruction under a maximum-likelihood frame-work. Only sites with high-confidence reconstructions (posterior probability ≥ 0.60) were retained, and terminal branches with *d_S_ >* 1 were excluded due to potential saturation (Supp. Table S6). For each gene–species combination, deviations from neutrality were assessed using Fisher’s exact test on contingency tables of nonsynonymous and synonymous polymorphism and divergence, and confidence intervals for NI were estimated from the corresponding odds ratios; *P*-values were adjusted for multiple testing using the Holm method. NI estimates spanned a wide range (median = 0.43; range = 0.02–22.6) and were highly variable, with wide and often unbounded confidence intervals, reflecting small counts and unstable denominators. Consistent with this, none of the comparisons were significant after multiple-testing correction (Holm-adjusted *P* = 1), indicating limited statistical power. In addition, underestimation of synonymous divergence due to saturation and posterior-probability filtering may bias NI values downward, further complicating interpretation. Thus, NI alone does not provide reliable evidence for positive selection on specific genes or branches.

Given these limitations, we next tested for episodic positive selection using likelihood-based models that allow some sites and lineages to evolve under positive selection while others evolve neutrally or under purifying selection, enabling us to distinguish adaptive changes from relaxed constraint. We first used site models (M8 vs. M8a), which test whether a subset of codons has *ω >* 1. Model M8 allows such sites, while M8a does not; if M8 fits significantly better, it suggests positive selection across the phylogeny (Yang et al., 2000). We did not find evidence for widespread positive selection. This is expected because positive selection is usually weak, occurs at only a few sites, and is often difficult to detect due to strong purifying selection, short alignments, low divergence, and multiple-testing correction.

We next applied branch-site models that allow *ω* to vary among sites and lineages, enabling tests for positive selection along specific foreground branches. Statistical sup-port was evaluated using likelihood ratio tests (LRTs), which test whether a model with positive selection fits better than a null model without it, and Akaike Information Criterion (AIC), which identifies the better-supported model while accounting for differences in complexity; *P*-values were Holm-adjusted. These analyses revealed a small amount of lineage-specific evidence of positive selection, with the strongest signals in *cox1* and *nadh5* (Table 2). In *P. calkinsi* and *P. caudatum*, *cox1* demonstrated strong support, and several codons were suggested to be under positive selection, which is consistent with episodic adaptation. A few genes (such as *rpL2*, *nadh2*) displayed marginal evidence below a strict *α* = 0.01 threshold, whereas *nadh5* displayed additional but weaker signals in *P. caudatum* and *P. multimicronucleatum*.

**Table 2.** Branch-site tests identifying lineage-specific positive selection in mitochondrial genes. Foreground branches correspond to the lineage on which selection was tested. Statistical support was evaluated using likelihood ratio tests (LRTs) and Akaike Information Criterion (AIC). Adjusted *P*-values were obtained using the Holm correction. Sites were identified using Bayes Empirical Bayes (BEB) analysis.

| Gene | Foreground lineage | Adj. $P$-value (Holm) | Rel. support (AIC) | Positively selected codons |
| --- | --- | --- | --- | --- |
| *cox1* | *P. caudatum* | $1.50 \times 10^{-7}$ | $7.65 \times 10^{-9}$ | 183S, 266S, 595C, 660G |
| *cox1* | *P. calkinsi* | $5.60 \times 10^{-7}$ | $2.76 \times 10^{-8}$ | 191P, 201S, 203F, 249V, 521G |
| *nadh5* | *P. caudatum* | $4.05 \times 10^{-5}$ | $1.78 \times 10^{-6}$ | 151A, 356N, 485L |
| *nadh5* | *P. multimicronucleatum* | $4.98 \times 10^{-5}$ | $2.18 \times 10^{-6}$ | 40S, 42I, 59L |
| *rpL2* | *P. multimicronucleatum* | $1.62 \times 10^{-3}$ | $6.25 \times 10^{-5}$ | 34M, 35P, 185I |
| *nadh2* | *P. multimicronucleatum* | $5.46 \times 10^{-3}$ | $2.01 \times 10^{-4}$ | 5L, 116K |

Structural predictions using DeepTMHMM indicate that candidate sites are predominantly located in loop regions that are exposed to the mitochondrial matrix or intermembrane space, rather than in transmembrane helices embedded within the lipid bilayer, which are subject to strong structural constraints due to the hydrophobic membrane environment (von Heijne, 2007). Overall, adaptive evolution in mitochondrial genes is rare and lineage-specific. Although *ymf* genes show relatively elevated *d_N_ /d_S_* and *π_N_ /π_S_*, both remain below 1, and the lack of branch-site support is consistent with weaker purifying selection rather than positive selection.

### Comparable effective population sizes in mitochondrial and nuclear genomes

We used synonymous diversity (*π_S_*) and divergence (*d_S_*) to compare the effective population sizes of the mitochondria and nuclei (Supp. Table S8). Under neutral expectations, *π_S_* = 2*N_e_µ* for mitochondria and *π_S_*= 4*N_e_µ* for the nucleus. Assuming that the divergence time (*t*) for nuclear and mitochondrial genomes is equal, *µ* can be estimated as proportional to *d_S_*. Synonymous divergence scales with mutation rate and time as *d_S_* ≈ 2*µt*. Accordingly, mitochondrial *d_S_* is consistently greater than nuclear *d_S_* in all species, with 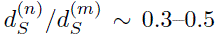 (Supp. Table 10), suggesting that mitochondrial mutation rates are around two–three times higher. The relative effective population size was thus calculated as follows:

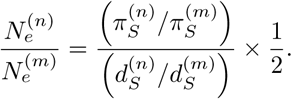

To avoid saturation effects, highly divergent species (*d_S_ >* 1; *P. caudatum*, *P. multimicronucleatum*, *P. bursaria*) were excluded.

The proportion of nuclear to mitochondrial effective population size 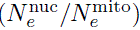 is 1.03 in *P. biaurelia*, whereas it drops slightly to about 0.90 in *P. primaurelia*, suggesting both are nearly matched. A noticeable shift appears in *P. tetraurelia*, where the value falls to around 0.53. By comparison, *P. sexaurelia* has the lowest ratio, which is approximately 0.10, indicating a more substantial mitochondrial *N_e_*, aligning with elevated synonymous diversity. Less restricted cytoplasmic inheritance and weak mitochondrial bottlenecks in *Paramecium* may help maintain mitochondrial *N_e_*at levels comparable to or higher than those of the nucleus (Adoutte and Beisson 1972; Adoutte et al. 1979; Lynch 2006). Overall, mitochondrial *N_e_* is comparable to or greater than nuclear *N_e_*, with variation across species.

### Selection in Mitochondrial and Nuclear-Encoded Functional Groups

Mitochondrial genomes appear to maintain effective population sizes similar to those of nuclear genomes despite limited evidence for recombination; yet, it is unclear how selection operates in the presence of both weak recombination and relatively large population sizes. To test this, we compared polymorphism (*π_N_ /π_S_*) and divergence (*dN/dS*) between mitochondrial and nuclear genes in the same functional classes, namely electron transport chain and ribosomal protein genes, using orthologs identified with OMA v2.6.0 (Altenhoff et al., 2019). The dataset includes nuclear-encoded electron transport chain (ETC) genes (∼66) and ribosomal proteins (∼83) (Fig. 4D; Supp. Table S9). For ETC genes, mitochondrial median *π_N_ /π_S_* (0.0605) and *dN/dS* (0.0436) are slightly lower than nuclear estimates (0.0666 and 0.0571), indicating comparable or slightly stronger purifying selection in mitochondria. For ribosomal proteins, mitochondrial median *π_N_ /π_S_* (0.0560) is also slightly lower than nuclear estimates (0.0642), whereas mitochondrial *dN/dS* is higher (0.0642 vs. 0.0445), consistent with broadly similar levels of constraint but some divergence in selective pressures between genomes.

Partitioning ribosomal proteins by subunit reveals contrasting patterns. In the nucleus, large-subunit (LSU) genes show higher median *π_N_ /π_S_* than small-subunit (SSU) genes (0.0709 vs. 0.0596), with nearly identical *dN/dS* values (0.0444 vs. 0.0448). In contrast, mitochondrial ribosomal proteins show the opposite trend, with SSU genes exhibiting higher *π_N_ /π_S_* (0.0766 vs. 0.0357) and *dN/dS* (0.0737 vs. 0.0570) than LSU genes. These opposing patterns indicate differential constraint across ribosomal subunits in nuclear versus mitochondrial genomes. Overall, these results are consistent with previous findings that mitochondrial genes do not show uniformly reduced efficacy of purifying selection and can exhibit comparable or stronger constraint depending on the gene set (Johri et al. 2019).

## Discussion

The mitochondrial genomes of the worldwide-studied species of *Paramecium* exhibit both considerable cryptic diversity and lineage-specific divergence in addition to conserved core gene content. Mitochondrial genes exhibit purifying selection similar to or stronger than the nuclear genomes despite the absence of recombination, consistent with mitochondrial effective population sizes that are not reduced relative to the nuclear genomes.

In contrast to the *P. aurelia* complex, mitochondrial genome of *P. bursaria* is considerably larger with apparent gaps in gene organization. These regions have unannotated ORFs within those gaps and are consistently found among isolates,indicating that they are actual genomic characteristics rather than nonfunctional intergenic regions. Many of these ORFs in this outgroup lineage exhibit inconsistent and weak similarity to ciliate-specific *ymf* genes, indicating that they are substantially divergent homologs whose sequence similarity is no longer easily detectable. These genes have higher polymorphism than core mitochondrial genes, indicating they may be able to tolerate a greater accumulation of mutations throughout evolutionary time.

Despite this lineage-specific expansion, the organization of the mitochondrial genome is generally preserved throughout *Paramecium*, with a stable core set of ribosomal and oxidative phosphorylation genes maintained across species. This tendency is consistent with strong functional constraints on energy metabolism that limit variation in gene content and organization even at large evolutionary distances. Similar levels of conservation have been reported in other ciliates, such as *Euplotes* and *Tetrahymena*, whose mitochondrial genomes maintain a conserved complement of core genes despite divergence in lineage-specific regions (Brunk et al., 2003; de Graaf et al., 2009; Burger et al., 2000). The essential role that oxidative phosphorylation plays in the generation of ATP, where even little disturbances can have a significant influence on fitness, is probably why these core genes have persisted. Because these organisms share the same natural environment, these restrictions might be more obvious. Effective respiratory metabolism is necessary for the free-living aquatic ciliates *Paramecium*, *Euplotes*, and *Tetrahymena* to grow and survive in oxygenated settings.

The process of mitochondrial inheritance and propagation is probably responsible for the maintenance of strong purifying selection in *Paramecium* mitochondrial genomes. *Paramecium* does not exhibit strict uniparental inheritance, unlike metazoans, in which mitochondria are passed down only through the maternal lineage and undergo a noticeable bottleneck during oogenesis. During conjugation, cytoplasmic exchange is limited but not absent, whereas autogamy involves self-fertilization without cytoplasmic mixing, thereby preserving existing mitochondrial populations. In both cases, cells maintain large numbers of mitochondria, and no severe bottleneck comparable to that in metazoans is imposed (Adoutte & Beisson, 1972). As a result, mitochondrial genomes are propagated at high copy number across generations and experience relatively weak reductions in effective population size. These features are expected to maintain relatively large mitochondrial *N_e_*, thereby enhancing the efficacy of purifying selection despite the absence of recombination (Lynch & Blanchard, 1998). This contrasts with metazoan systems, where reduced mitochondrial *N_e_* is thought to contribute to the accumulation of slightly deleterious mutations (Stewart & Chinnery, 2015).

These results demonstrate the importance of mitochondrial genomics for understanding diversification of microbial eukaryotes and provide a framework for linking the evolution of the mitochondrial genome to adaptability and speciation in ciliates.

## Methods

### Sample Collection and Culturing

Environmental water samples were gathered from freshwater habitats in various geographical locations that had loamy soils and stagnant water. Samples were collected from several locations and time intervals in order to capture independent isolates. Shortly after collection, the samples were separated into different containers to reduce clonality. To encourage microbial development, small amounts of rice were added to the samples. *Paramecium* isolates were obtained within a few days of collection by micropipette isolation and serial washing to reduce contamination. Individual cells were transferred to culture media (Dryl’s solution, wheatgrass, or lettuce medium) and maintained with Klebsiella aerogenes (ATCC 35028) as a food source. Cultures were grown under standard conditions and expanded for downstream DNA extraction.

### DNA isolation, sequencing, assembly, and species identification

We extracted genomic DNA using standard kits (MasterPure or MagMax) and sequenced it on an Illumina with 150 bp paired-end reads. Before combining samples, we checked the quality of the libraries using the NEBNext Ultra II FS kit while keeping PCR to a minimum (just 6–7 cycles) to avoid over-amplifying anything. Reads were trimmed using Trimmomatic (Bolger et al., 2014), mapped to nuclear reference genomes to remove nuclear DNA. We also ran Kraken to screen out bacterial reads so contamination wouldn’t carry through (Wood & Salzberg, 2014). With the remaining mitochondrial-enriched reads, we assembled genomes using SPAdes meta (v3.15.5) (Bankevich et al., 2012). To identify the species, we used whole genome to BLAST searches against ParameciumDB (Altschul et al., 1990; Arnaiz et al., 2019). Assembly quality was assessed using coverage thresholds for nuclear genomes (≥60% breadth, ≥20–30× depth) and mitochondrial criteria, including ≥95% *cox1* identity and ≥90% read coverage across the scaffold.

### Mitochondrial genome annotation

We found open reading frames (ORFs) by scanning each assembly in all six reading frames to identify mitochondrial protein-coding genes. ORFs were translated using the *Paramecium* mitochondrial genetic code (NCBI translation table 4), where UGA encodes tryptophan and other start codons (such as UUG) (Arnaiz et al., 2019). The longest ORF was kept for each ORF cluster, utilizing BLAST to compare it to a reference set of 46 mitochondrial proteins (Johri et al., 2019). The best-scoring match was kept after hits were filtered using an E-value threshold of *<* 0.001 and ≥30% coverage to guarantee one-to-one gene assignment. Infernal with Rfam covariance models was used to annotate ribosomal RNA genes, and tRNAscan-SE v2 in organellar mode was used to identify transfer RNAs (Nawrocki & Eddy, 2013; Chan & Lowe, 2019).

### Phylogenetic analysis

Amino acid sequences from annotated mitochondrial protein-coding genes were aligned across species using MAFFT v7.525 (Katoh & Standley, 2013).BMGE was used to eliminate poorly aligned regions: –t AA –m BLOSUM62 –g 0.5 –h 0:0.5 –w 3 –b 5; (Criscuolo & Gribaldo, 2010). The filtered alignments were then concatenated into a supermatrix. A maximum-likelihood tree was inferred using RAxML-NG under the LG+G model with 100 bootstrap replicates: raxml-ng-mpi ––all ––model LG+G ––data-type AA ––tree rand{10} ––bs-trees 100; (Kozlov et al., 2019). The same alignment was also examined using 1,000 ultrafast bootstraps and 1,000 SH-aLRT tests as an independent check using IQ-TREE2 (Minh et al., 2020). Individual gene trees were inferred using RAxML-NG with the same model and bootstrap parameters in order to evaluate gene-level support. A majority-vote method was used to assess important backbone relationships across gene trees, and the results were summarized into a gene-tree consensus table (Supp. Table 4).

### Mitochondrial genome structure and composition analysis

We annotated mitochondrial protein-coding genes, tRNAs, and rRNAs in *Paramecium* and plotted gene maps in R with ggplot2/gggenes (Wickham, 2016; Wilkins & Kurtz, 2025). Gene length was measured using coding-sequence coordinates from mitochondrial genes consistently identified across all genomes. Gene lengths were normalized by dividing species-specific values by the median length of each ortholog across all species. Gene-level estimations were calculated for all identified protein-coding genes, and mitochondrial GC content was simultaneously measured at the whole-genome and gene-wise levels. Comparisons were performed between the *P. aurelia* complex and remaining outgroup species (*P. bursaria*, *P. tribursaria*, *P. caudatum*, *P. multimicronucleatum*, *P. fokini*, and *P. calkinsi*).

### Nucleotide Diversity Estimation Across Mitochondrial Site Classes

Nucleotide diversity (*π*) was calculated for mitochondrial genomes from multiple *Paramecium* species in order to quantify patterns of variation among various functional site classes. For protein-coding genes, codon-aware multiple sequence alignments were constructed, and gene-specific degeneracy maps were used to categorize each nucleotide location by degeneracy (0, 2, 3, or 4-fold). Per-site nucleotide diversity was computed using the average number of pairwise differences between two sequences as:

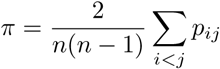

where *n* is the number of sequences at a site that have valid nucleotide calls and *p_ij_*counts the number of pairs of sequences that differ at the site. Sites containing gaps or ambiguous bases were excluded. Site-weighted mean *π* estimates for each species were then calculated by averaging site-wise *π* values across all sites within each codon degeneracy class (Bruen et al., 2006). Mitochondrial tRNA, rRNA, and intergenic regions were aligned independently using MAFFT. Per-site nucleotide diversity *π* was computed for each alignment. The site-weighted mean nucleotide diversity estimates for RNA genes and intergenic regions were then calculated by adding the site-specific *π* values across all regions and dividing by the total length of such regions within each species.

### Tests for Recombination

Recombination in mitochondrial protein-coding genes was assessed using PhiPack (Bruen et al. 2006). For each species, codon-aligned nucleotide sequences were analyzed gene-wise using a 100 bp sliding window and 1,000 permutations. Four complementary tests were applied for each gene–species combination: the PHI test (permutation and normal approximation), MaxChi2, which detects shifts in site compatibility across alignments, and the Neighbor Similarity Score (NSS), which measures clustering of compatible sites. These methods detect recombination through nonrandom associations among polymorphic sites, conceptually similar to the four-gamete (four-allele) criterion under the infinite-sites model, but without relying solely on distance-dependent decay of linkage disequilibrium. Results were summarized in gene × species matrices and visualized as heatmaps, with outcomes classified as significant (*P <* 0.05), non-significant, or missing.

Recombination was analyzed using the pairwise module of LDhat v2.2 (McVean et al., 2002, 2004). Analyses were performed per species using mitochondrial SNP datasets restricted to homozygous, biallelic sites with known ancestral states. Both crossover and gene-conversion models (mean tract length = 500 bp) were tested, with two values of *θ* (4*N_e_µ*) and a recombination grid up to 4*N_e_r* = 100 (101 points). Evidence for recombination was assessed using likelihood ratio tests, the *G*_4_ statistic, and permutation tests (1,000 replicates) of linkage disequilibrium decay with distance (*r*^2^, *D^′^*). Recombination rates were estimated using the estimators of Wakeley (1997) and Hudson (finite-sites model).

### Within-species *π_N_ /π_S_* estimates across mitochondrial genes

Nonsynonymous and synonymous nucleotide diversity (*π_N_* and *π_S_*) within species were evaluated using codon-aligned mitochondrial protein-coding sequences for each species. Each gene’s consensus coding sequence was created using a ≥ 60% majority rule per nucleotide; codons with gaps, ambiguous bases, or stop codons were eliminated. By counting all single-nucleotide substitutions at each codon position, the total number of synonymous (S-sites) and nonsynonymous (N-sites) sites per gene was ascertained (translation table 4). All pairwise sequence comparisons among species were then carried out codon by codon, and any differences were categorized as either synonymous or nonsynonymous. The mean per-site diversity estimates (*π_S_*and *π_N_*) for each gene were obtained by dividing the mean numbers of synonymous and nonsynonymous differences across all sequence pairs by S-sites and N-sites, respectively. The ratio (*π_N_ /π_S_*) was utilized as a measure of within-species selective constraint (Nei & Gojobori, 1986; Nei & Li, 1979).

### Between-species dN/dS and inference of site-wise selection coefficients across mitochondrial genes

The site model M8 (beta + *ω*) implemented in codeml from PAML (Yang, 2007) was used to quantify selective pressures on mitochondrial protein-coding genes. Codon alignments were examined under model = 0, NSsites = 8, CodonFreq = 2, and icode = 4. Model M8 allows *ω* (*d_N_ /d_S_*) to vary among sites, with most sites following a beta distribution (0 *< ω <* 1) and an additional class permitting *ω >* 1 (Yang et al. 2000). Mean *ω* values were used to summarize gene-level selective constraint.

To interpret these estimates in a population-genetic framework, *ω* distributions were transformed into scaled selection coefficients (*S*) using 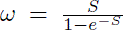 (Halpern & Bruno, 1998; Yang & Nielsen, 2008). Analyses were restricted to genes with the majority of sites assigned to the beta-distributed class under the M8 model (*p*_0_ *>* 0.999). Continuous distributions of the scaled selection coefficient (*S*; −20 ≤ *S* ≤ 5) were generated, and the proportion of sites with *S >* 1 and the modal value for *S <* 0 were used to summarize positive and purifying selection, respectively.

### Neutrality Index

Under a single-ratio codon model (model = 0, NSsites = 0) (Yang 2007), ancestral sequences for every mitochondrial gene were rebuilt using codeml in PAML, with ancestral states inferred at internal nodes (RateAncestor = 2). By comparing current sequences to reconstructed ancestors and excluding sites with gaps, stop codons, or ambiguous bases, branch-specific nonsynonymous (*d_N_*) and synonymous (*d_S_*) substitution rates (number of substitutions per sites) were found. To minimize uncertainty, analyses were limited to sites where the posterior probability of site-specific reconstruction ≥ 0.60. The same filtered sites were used to construct polymorphism-based diversity (*π_N_*, *π_S_*) and divergence estimates (*d_N_*, *d_S_*), removing gene–species pairs with zero counts or *d_S_* ≥ 1.The neutrality index was calculated as NI = (*π_N_ /π_S_*)*/*(*d_N_ /d_S_*) (McDonald & Kreitman, 1991; Rand & Kann, 1996). Fisher’s exact test was used to evaluate statistical significance on 2 × 2 contingency tables, and the Holm technique was used to adjust the *P*-values (Holm, 1979).

### Tests for positive selection using site and branch-site models

Site and branch-site codon models implemented in codeml from PAML were used to test for positive selection (Yang, 2007). For every gene, codon alignments and fixed phylogenetic trees were utilized as input. To find widespread positive selection across lineages, site models M8 and M8a (model = 0, NSsites = 8) were examined. Both models permit *ω* (*d_N_ /d_S_*) to fluctuate between sites in accordance with a beta distribution; however, M8a restricts this class to *ω* = 1, while M8 includes an extra site class where *ω >* 1 (Yang et al. 2000). Likelihood ratio tests (LRTs) were used to assess model fit, and the test statistic 2Δ*ℓ* = 2(ln *L*_M8_ − ln *L*_M8a_) was compared to a *χ*^2^ distribution with one degree of freedom. The Holm technique (Holm 1979) was used to adjust *P*-values for multiple testing.

To detect episodic positive selection, branch-site Model A was applied (model = 2, NSsites = 2), allowing a subset of sites to evolve with *ω >* 1 on specified foreground branches (Zhang et al. 2005). Two foreground configurations were tested: each terminal branch individually, and all *P. aurelia* lineages jointly.Model A was compared to a null model with *ω* = 1 in each case using LRTs. A *χ*^2^ distribution with one degree of freedom was used to test the test statistic 2Δ*ℓ* = 2(ln *L*_ModelA_ − ln *L*_null_). Relative Akaike weights were calculated and model support was further assessed using the Akaike Information Criterion (AIC), which is defined as AIC = 2*k* − 2 ln *L*. Codon sites under positive selection were found using Bayes Empirical Bayes (BEB) analysis (Yang et al. 2005).

### Identification of Nuclear-Encoded Orthologs and Functional Gene Sets for Selection Analyses

OMA v2.6.0 was used to identify orthologous genes among *Paramecium* species (Altenhoff et al., 2019), producing 39,383 orthologous groups at the root level. To define nuclear-encoded electron transport chain (ETC) genes, we curated components of mitochondrial respiratory complexes based on structural characterization in *Tetrahymena* (Zhou et al., 2022) and identified homologs in *P. caudatum* using sequence similarity searches with an E-value cutoff of 1 × 10*^−^*^3^ (0.001). A final set of approximately 66 ETC genes was obtained by validating the presence of candidate genes in *P. tetraurelia* and using OMA hierarchical orthologous groups to recover orthologs across species. For ribosomal proteins, we used structural annotations of the mitochondrial large and small ribosomal subunits from yeast (Amunts et al., 2014; Desai et al., 2017) to identify conserved components, followed by homolog identification in *P. bursaria* and *P. caudatum* using the same E-value threshold (1 × 10*^−^*^3^), and ortholog retrieval across species via OMA, yielding approximately 83 ribosomal protein (RP) genes. Gene-level coding sequences were extracted and translated using the ciliate nuclear genetic code (genetic code 6), and codon alignments were used to estimate within-species polymorphism (*π_N_ /π_S_*) and between-species divergence (*d_N_ /d_S_*) for both functional groups.

## Acknowlegements

This research was facilitated by the Biodesign Centre for Mechanisms of Evolution, Arizona State University, USA. This work was supported by the National Science Foundation (DEB-1927159 and DBI-2119963), the National Institutes of Health (R35-GM122566-08), and Grant 735927 from the Moore–Simons Project, awarded to Michael Lynch, and by National Science Foundation grant IOS-1838098, awarded to John P. DeLong and Kristi L. Montooth.

## Supplementary Information

**Table S1:** Sample names and collection localities for all mitochondrial genome assemblies analyzed in this study.

| Species | Sample | Collection |
| --- | --- | --- |
| *P. biaurelia* | 11_IV_2 | USA |
| *P. biaurelia* | 12-I-16 | Bloomington, Indiana, USA |
| *P. biaurelia* | 294 | Germany |
| *P. biaurelia* | 36-I-2 | USA |
| *P. biaurelia* | 36_I_1 | USA |
| *P. biaurelia* | 4-IV-2 | USA |
| *P. biaurelia* | 4_IV_3 | USA |
| *P. biaurelia* | A1-4 | Germany |
| *P. biaurelia* | AL7-10 | Ayer Lake, Arizona |
| *P. biaurelia* | AL7-25 | Ayer Lake, Arizona |
| *P. biaurelia* | AL7-27_B | Ayer Lake, Arizona |
| *P. biaurelia* | BA1C_2 | Basha Kill Marsh, New York |
| *P. biaurelia* | BA2A_1 | Basha Kill Marsh, New York |
| *P. biaurelia* | BA4C_1 | Basha Kill Marsh, New York |
| *P. biaurelia* | CL12-1 | Canyon Lake, Arizona |
| *P. biaurelia* | CL12-3 | Canyon Lake, Arizona |
| *P. biaurelia* | CL12-8 | Canyon Lake, Arizona |
| *P. biaurelia* | CL5-12 | Canyon Lake, Arizona |
| *P. biaurelia* | DP1G_2 | Duck Pond, New York |
| *P. biaurelia* | G10 | USA |
| *P. biaurelia* | G11 | USA |
| *P. biaurelia* | G7 | USA |
| *P. biaurelia* | G8 | USA |
| *P. biaurelia* | G9 | USA |
| *P. biaurelia* | M2Hc | Germany |
| *P. biaurelia* | Opa | Opatowice, Poland |
| *P. biaurelia* | P2a.45 | Israel |
| *P. biaurelia* | P_2a-44 | Spain |
| *P. biaurelia* | RL12-10 | Roosevelt Lake, Arizona |
| *P. biaurelia* | WCL3-10 | Woods Canyon, Arizona |
| *P. biaurelia* | YK2H_1 | Yanketown Pond, New York |
| *P. biaurelia* | biaurelia_V1_4 | Arizona |
| *P. bursaria* | AZ1 | Arizona |
| *P. bursaria* | AZ4 | Arizona |
| *P. bursaria* | BB1 | Baden-Baden, Germany |
| *P. bursaria* | C173 | Germany |
| *P. bursaria* | MW_45 | Marsh Wren Marsh, Nebraska |
| *P. bursaria* | P1-12B | Ramsey Canyon Preserve, Arizona |
| *P. bursaria* | P1-14B | Ramsey Canyon Preserve, Arizona |
| *P. bursaria* | P1-16_B | Ramsey Canyon Preserve, Arizona |
| *P. bursaria* | P1-19B | Ramsey Canyon Preserve, Arizona |
| *P. bursaria* | P1-20_B | Ramsey Canyon Preserve, Arizona |
| *P. bursaria* | P1-21_B | Ramsey Canyon Preserve, Arizona |
| *P. bursaria* | P1-3a_B | Ramsey Canyon Preserve, Arizona |
| *P. bursaria* | P1-7a_(B) | Ramsey Canyon Preserve, Arizona |
| *P. bursaria* | P1-8a(B) | Ramsey Canyon Preserve, Arizona |
| *P. bursaria* | P2-16 | Ramsey Canyon Preserve, Arizona |
| *P. bursaria* | P2-20_(B) | Ramsey Canyon Preserve, Arizona |
| *P. bursaria* | P2-4_(B) | Ramsey Canyon Preserve, Arizona |
| *P. bursaria* | P2-6_(B) | Ramsey Canyon Preserve, Arizona |
| *P. bursaria* | P2-7B | Ramsey Canyon Preserve, Arizona |
| *P. bursaria* | P2-9_(B) | Ramsey Canyon Preserve, Arizona |
| *P. bursaria* | P3-16aB | Ramsey Canyon Preserve, Arizona |
| *P. bursaria* | P3-17a | Ramsey Canyon Preserve, Arizona |
| *P. bursaria* | P3-17b | Ramsey Canyon Preserve, Arizona |
| *P. bursaria* | P3-19_(B) | Ramsey Canyon Preserve, Arizona |
| *P. bursaria* | P3-3b_(B)2nd | Ramsey Canyon Preserve, Arizona |
| *P. bursaria* | P3-5a(B) | Ramsey Canyon Preserve, Arizona |
| *P. bursaria* | SC43B | Spring Creek Pond, Nebraska |
| *P. bursaria* | SC_15_B | Spring Creek Pond, Nebraska |
| *P. bursaria* | SC_17B | Spring Creek Pond, Nebraska |
| *P. bursaria* | SC_22-13B | Spring Creek Pond, Nebraska |
| *P. bursaria* | SC_22-29B | Spring Creek Pond, Nebraska |
| *P. bursaria* | SC_38_B | Spring Creek Pond, Nebraska |
| *P. bursaria* | SC_3_B | Spring Creek Pond, Nebraska |
| *P. bursaria* | SC_4B | Spring Creek Pond, Nebraska |
| *P. caudatum* | 6.II.4 | Monroe Lake, Indiana, USA |
| *P. caudatum* | C082 | USA |
| *P. caudatum* | C139 | Japan |
| *P. caudatum* | C171 | Greece |
| *P. caudatum* | CO30 | Austria |
| *P. caudatum* | HC10-10 | Horton Creek, Arizona |
| *P. caudatum* | HC6-13 | Horton Creek, Arizona |
| *P. caudatum* | HC6-15 | Horton Creek, Arizona |
| *P. caudatum* | HC6-16 | Horton Creek, Arizona |
| *P. caudatum* | HC6-5 | Horton Creek, Arizona |
| *P. caudatum* | HC6-7 | Horton Creek, Arizona |
| *P. caudatum* | HC7-9 | Horton Creek, Arizona |
| *P. caudatum* | HC8-6a | Horton Creek, Arizona |
| *P. caudatum* | HC8-7 | Horton Creek, Arizona |
| *P. caudatum* | HC9-9a_Large | Horton Creek, Arizona |
| *P. caudatum* | LP3-8 | Lake Pleasant, Arizona |
| *P. caudatum* | LP6-1 | Lake Pleasant, Arizona |
| *P. caudatum* | LP6-3 | Lake Pleasant, Arizona |
| *P. caudatum* | LP6-10 | Lake Pleasant, Arizona |
| *P. caudatum* | LP7-3 | Lake Pleasant, Arizona |
| *P. caudatum* | LP7-5a | Lake Pleasant, Arizona |
| *P. caudatum* | LP7-10 | Lake Pleasant, Arizona |
| *P. caudatum* | LP7-12 | Lake Pleasant, Arizona |
| *P. caudatum* | LP7-14 | Lake Pleasant, Arizona |
| *P. caudatum* | LP8-12 | Lake Pleasant, Arizona |
| *P. caudatum* | NF23 | Fukushima Daiichi Power Plant, Japan |
| *P. caudatum* | NF4 | Fukushima Daiichi Power Plant, Japan |
| *P. caudatum* | NF_13 | Fukushima Daiichi Power Plant, Japan |
| *P. caudatum* | NF_20 | Fukushima Daiichi Power Plant, Japan |
| *P. caudatum* | NF_3 | Fukushima Daiichi Power Plant, Japan |
| *P. caudatum* | RL15-37 | Roosevelt Lake, Arizona |
| *P. caudatum* | RL15-61 | Roosevelt Lake, Arizona |
| *P. caudatum* | SC22-7 | Spring Creek Pond, Nebraska |
| *P. caudatum* | SC22-16 | Spring Creek Pond, Nebraska |
| *P. caudatum* | SC22-20 | Spring Creek Pond, Nebraska |
| *P. caudatum* | SC22-22 | Spring Creek Pond, Nebraska |
| *P. caudatum* | SC22-27 | Spring Creek Pond, Nebraska |
| *P. caudatum* | SC22-28 | Spring Creek Pond, Nebraska |
| *P. caudatum* | SC22-36 | Spring Creek Pond, Nebraska |
| *P. caudatum* | SF_3 | Fukushima Daiichi Power Plant, Japan |
| *P. caudatum* | SF_4 | Fukushima Daiichi Power Plant, Japan |
| *P. caudatum* | SF_5 | Fukushima Daiichi Power Plant, Japan |
| *P. caudatum* | SF_6 | Fukushima Daiichi Power Plant, Japan |
| *P. caudatum* | SF_9 | Fukushima Daiichi Power Plant, Japan |
| *P. caudatum* | WC1-6a | Woods Canyon, Arizona |
| *P. caudatum* | WCL2-11 | Woods Canyon, Arizona |
| *P. caudatum* | WCL3-6 | Woods Canyon, Arizona |
| *P. caudatum* | WCL5-3 | Woods Canyon, Arizona |
| *P. caudatum* | WCL5-4 | Woods Canyon, Arizona |
| *P. caudatum* | WCL8-15 | Woods Canyon, Arizona |
| *P. caudatum* | WCL_2-4 | Woods Canyon, Arizona |
| *P. caudatum* | WCL_2-5 | Woods Canyon, Arizona |
| *P. caudatum* | WCL_2-8 | Woods Canyon, Arizona |
| *P. caudatum* | WCL_2-10 | Woods Canyon, Arizona |
| *P. caudatum* | WCL_2-12 | Woods Canyon, Arizona |
| *P. caudatum* | WCL_2-15 | Woods Canyon, Arizona |
| *P. caudatum* | WL1-1C | Watson Lake, Arizona |
| *P. caudatum* | WL2-3B | Watson Lake, Arizona |
| *P. caudatum* | WL5-2C | Watson Lake, Arizona |
| *P. caudatum* | WL5-6B | Watson Lake, Arizona |
| *P. caudatum* | WL5-7B | Watson Lake, Arizona |
| *P. caudatum* | caudatum_43c3d | Arizona |
| *P. caudatum* | caudatum_C026 | Baton Rouge, USA |
| *P. caudatum* | caudatum_C083 | Bloomington, USA |
| *P. caudatum* | caudatum_C104 | Spain |
| *P. caudatum* | caudatum_GBE | Arizona |
| *P. fokini* | MW23 | Marsh Wren Marsh, Nebraska |
| *P. fokini* | MW24 | Marsh Wren Marsh, Nebraska |
| *P. fokini* | MW_11 | Marsh Wren Marsh, Nebraska |
| *P. fokini* | MW_12 | Marsh Wren Marsh, Nebraska |
| *P. fokini* | MW_14 | Marsh Wren Marsh, Nebraska |
| *P. fokini* | MW_15 | Marsh Wren Marsh, Nebraska |
| *P. fokini* | MW_18 | Marsh Wren Marsh, Nebraska |
| *P. fokini* | MW_20 | Marsh Wren Marsh, Nebraska |
| *P. fokini* | MW_21 | Marsh Wren Marsh, Nebraska |
| *P. fokini* | MW_22 | Marsh Wren Marsh, Nebraska |
| *P. fokini* | MW_25 | Marsh Wren Marsh, Nebraska |
| *P. fokini* | MW_26 | Marsh Wren Marsh, Nebraska |
| *P. fokini* | MW_27 | Marsh Wren Marsh, Nebraska |
| *P. fokini* | MW_30 | Marsh Wren Marsh, Nebraska |
| *P. fokini* | MW_31 | Marsh Wren Marsh, Nebraska |
| *P. fokini* | MW_32 | Marsh Wren Marsh, Nebraska |
| *P. fokini* | MW_33 | Marsh Wren Marsh, Nebraska |
| *P. fokini* | MW_34 | Marsh Wren Marsh, Nebraska |
| *P. fokini* | MW_3 | Marsh Wren Marsh, Nebraska |
| *P. fokini* | MW_4 | Marsh Wren Marsh, Nebraska |
| *P. fokini* | MW_5 | Marsh Wren Marsh, Nebraska |
| *P. fokini* | MW_6 | Marsh Wren Marsh, Nebraska |
| *P. fokini* | MW_7 | Marsh Wren Marsh, Nebraska |
| *P. fokini* | MW_8 | Marsh Wren Marsh, Nebraska |
| *P. fokini* | NE1 | Marsh Wren Marsh, Nebraska |
| *P. fokini* | NE3 | Marsh Wren Marsh, Nebraska |
| *P. fokini* | NE4 | Marsh Wren Marsh, Nebraska |
| *P. fokini* | NE8 | Marsh Wren Marsh, Nebraska |
| *P. fokini* | NJH-QZ | China |
| *P. fokini* | NJH-SZ | China |
| *P. fokini* | NJH-WN | China |
| *P. fokini* | Pmmn_MO3C4 | Germany |
| *P. fokini* | SC22-13 | Spring Creek Pond, Nebraska |
| *P. fokini* | SC22-19 | Spring Creek Pond, Nebraska |
| *P. fokini* | SC22-25 | Spring Creek Pond, Nebraska |
| *P. fokini* | SC22-26 | Spring Creek Pond, Nebraska |
| *P. fokini* | SC_22-1 | Spring Creek Pond, Nebraska |
| *P. multimicronucleatum* | AL7-30 | Ayer Lake, Arizona |
| *P. multimicronucleatum* | MW_40 | Marsh Wren Marsh, Nebraska |
| *P. multimicronucleatum* | MW_42 | Marsh Wren Marsh, Nebraska |
| *P. multimicronucleatum* | NE5 | Marsh Wren Marsh, Nebraska |
| *P. multimicronucleatum* | PPP22-17 | Nebraska |
| *P. multimicronucleatum* | RL6B | Roosevelt Lake, Arizona |
| *P. multimicronucleatum* | SWL_1-25 | Wisconsin, Arizona |
| *P. multimicronucleatum* | multimicronucleatum_3I | Portugal |
| *P. quindecarelia* | CL11-12 | Canyon Lake, Arizona |
| *P. quindecarelia* | CL11-13 | Canyon Lake, Arizona |
| *P. quindecarelia* | CL11-14 | Canyon Lake, Arizona |
| *P. quindecarelia* | CL11-16 | Canyon Lake, Arizona |
| *P. quindecarelia* | CL11-19 | Canyon Lake, Arizona |
| *P. quindecarelia* | CL11-20 | Canyon Lake, Arizona |
| *P. quindecarelia* | CL11-24 | Canyon Lake, Arizona |
| *P. quindecarelia* | HC6-14 | Horton Creek, Arizona |
| *P. quindecarelia* | HC9-9b | Horton Creek, Arizona |
| *P. quindecarelia* | WCL_3-12 | Woods Canyon, Arizona |
| *P. primaurelia* | 14_II_2 | Lake Lemon, Indiana, USA |
| *P. primaurelia* | 3_III_1 | Bloomington, Indiana, USA |
| *P. primaurelia* | AL1-1 | Ayer Lake, Arizona |
| *P. primaurelia* | AL1-4 | Ayer Lake, Arizona |
| *P. primaurelia* | AL2-10 | Ayer Lake, Arizona |
| *P. primaurelia* | AL3-6 | Ayer Lake, Arizona |
| *P. primaurelia* | AL3-7 | Ayer Lake, Arizona |
| *P. primaurelia* | AL3-B | Ayer Lake, Arizona |
| *P. primaurelia* | AL3B | Ayer Lake, Arizona |
| *P. primaurelia* | AL4-A | Ayer Lake, Arizona |
| *P. primaurelia* | AL6A | Ayer Lake, Arizona |
| *P. primaurelia* | C124 | Germany |
| *P. primaurelia* | CAC1-14 | California |
| *P. primaurelia* | CAC1-16 | California |
| *P. primaurelia* | CAC1-17 | California |
| *P. primaurelia* | CAC1-18 | California |
| *P. primaurelia* | CAC1-21 | California |
| *P. primaurelia* | CAC1-22 | California |
| *P. primaurelia* | CAC1-23 | California |
| *P. primaurelia* | CAC1-25 | California |
| *P. primaurelia* | CAC1-2 | California |
| *P. primaurelia* | CAC1-3 | California |
| *P. primaurelia* | CAC1-9 | California |
| *P. primaurelia* | CL11-10 | Canyon Lake, Arizona |
| *P. primaurelia* | CL11-15 | Canyon Lake, Arizona |
| *P. primaurelia* | CL11-3 | Canyon Lake, Arizona |
| *P. primaurelia* | CL3-12 | Canyon Lake, Arizona |
| *P. primaurelia* | CL5-11 | Canyon Lake, Arizona |
| *P. primaurelia* | CL5-1 | Canyon Lake, Arizona |
| *P. primaurelia* | CL5-2 | Canyon Lake, Arizona |
| *P. primaurelia* | CL_1-14 | Canyon Lake, Arizona |
| *P. primaurelia* | CL_22-10 | Nebraska |
| *P. primaurelia* | CL_22-7 | Nebraska |
| *P. primaurelia* | Ecuador_A1-9_I | Ecuador |
| *P. primaurelia* | G5 | Istanbul, Turkey |
| *P. primaurelia* | Ista1_d.1_IV | Istanbul, Turkey |
| *P. primaurelia* | Istanbul-1a-III | Istanbul, Turkey |
| *P. primaurelia* | LP4-2b | Lake Pleasant, Arizona |
| *P. primaurelia* | LP7-1 | Lake Pleasant, Arizona |
| *P. primaurelia* | LP7-4 | Lake Pleasant, Arizona |
| *P. primaurelia* | LP8-13 | Lake Pleasant, Arizona |
| *P. primaurelia* | RL13-15b | Roosevelt Lake, Arizona |
| *P. primaurelia* | RL7B | Roosevelt Lake, Arizona |
| *P. primaurelia* | Rio_Lg_Jac.2_II | Brazil |
| *P. primaurelia* | SC22-12 | Spring Creek Pond, Nebraska |
| *P. primaurelia* | SC34B | Spring Creek Pond, Nebraska |
| *P. primaurelia* | YE_4 | USA |
| *P. primaurelia* | YE_5 | USA |
| *P. primaurelia* | YKKWold | Honsyu Island, Japan |
| *P. primaurelia* | primaurelia_AZ9-3 | Arizona |
| *P. quadecaurelia* | SL1-A | Arizona |
| *P. quadecaurelia* | SL5B | Arizona |
| *P. quadecaurelia* | quadecaurelia_NiA | USA |
| *P. sexaurelia* | 11_II_7 | USA |
| *P. sexaurelia* | A6-132 | Spain |
| *P. sexaurelia* | A6-133 | Spain |
| *P. sexaurelia* | A6-137 | Russia |
| *P. sexaurelia* | C131 | Ecuador |
| *P. sexaurelia* | Ecuad_6_II | Ecuador |
| *P. sexaurelia* | G3 | Japan |
| *P. sexaurelia* | Indo_1_9_II | Indonesia |
| *P. sexaurelia* | Indon_.1.7_I | Indonesia |
| *P. sexaurelia* | Indon_.12.5_I | Indonesia |
| *P. sexaurelia* | J2 | Indonesia |
| *P. sexaurelia* | Moz_13_C_IX | Mozambique |
| *P. sexaurelia* | P_6a-131 | Germany |
| *P. sexaurelia* | RL6-2 | Roosevelt Lake, Arizona |
| *P. sexaurelia* | sexaurelia_128 | Japan |
| *P. sexaurelia* | sexaurelia_130 | Ioannina Lake, Greece |
| *P. sexaurelia* | sexaurelia_133 | Seville, Spain |
| *P. sexaurelia* | sexaurelia_AZ8-4 | Arizona |
| *P. tetraurelia* | 116-K | Bloomington, Indiana, USA |
| *P. tetraurelia* | 169k | Morioka city, Japan |
| *P. tetraurelia* | 198_K_24C | Germany |
| *P. tetraurelia* | 290apo | Germany |
| *P. tetraurelia* | 298_KC+ | Germany |
| *P. tetraurelia* | 298_apo | Madrid, Spain |
| *P. tetraurelia* | 3_III_2 | USA |
| *P. tetraurelia* | 47 | Berkley, California, USA |
| *P. tetraurelia* | 47apo | Berkley, California, USA |
| *P. tetraurelia* | 51_apo | Spencer, Indiana, USA |
| *P. tetraurelia* | 98apo | Germany |
| *P. tetraurelia* | A30_K | Littlehampton, Australia |
| *P. tetraurelia* | A30cplus | Germany |
| *P. tetraurelia* | A4-116 | USA |
| *P. tetraurelia* | A4-291 | Australia |
| *P. tetraurelia* | A4-293 | Australia |
| *P. tetraurelia* | A4-298 | Panama |
| *P. tetraurelia* | Bapo | Krakow, Poland |
| *P. tetraurelia* | Brazil_P3 | Brazil |
| *P. tetraurelia* | G1 | Germany |
| *P. tetraurelia* | Istanbul_2_b_IV_3 | Istanbul, Turkey |
| *P. tetraurelia* | Moz_13_C_XI | Mozambique |
| *P. tetraurelia* | tetraurelia_32 | Arizona |
| *P. tetraurelia* | tetraurelia_51 | Spencer, USA |
| *P. tredecaurelia* | Pa_a_PFE_42 | Germany |
| *P. tredecaurelia* | tredecaurelia_209 | USA |
| *P. triaurelia* | CL12-12 | Canyon Lake, Arizona |
| *P. triaurelia* | CL12-18 | Canyon Lake, Arizona |
| *P. triaurelia* | CL12-27 | Canyon Lake, Arizona |
| *P. triaurelia* | CL_22-29 | Nebraska |
| *P. triaurelia* | CL_22-33 | Nebraska |
| *P. triaurelia* | CL_22-37 | Nebraska |
| *P. triaurelia* | CL_22-39 | Nebraska |
| *P. triaurelia* | LP2-1b | Lake Pleasant, Arizona |
| *P. triaurelia* | LP3-1 | Lake Pleasant, Arizona |
| *P. triaurelia* | LP3-6 | Lake Pleasant, Arizona |
| *P. triaurelia* | LP6-15 | Lake Pleasant, Arizona |
| *P. triaurelia* | LP7-5b | Lake Pleasant, Arizona |
| *P. triaurelia* | SA1E_1 | Great Sacandaga Lake, New York |
| *P. tribursaria* | MW_46 | Marsh Wren Marsh, Nebraska |
| *P. tribursaria* | RL13-_18a_B | Roosevelt Lake, Arizona |
| *P. tribursaria* | RL13-_4a_B | Roosevelt Lake, Arizona |
| *P. tribursaria* | RL14-15b-B | Roosevelt Lake, Arizona |
| *P. tribursaria* | RL14-_9b_B | Roosevelt Lake, Arizona |
| *P. tribursaria* | RL15-14b-B | Roosevelt Lake, Arizona |
| *P. tribursaria* | WC9-1b | Woods Canyon, Arizona |
| *P. tribursaria* | Yad1g1N | Honsyu Island, Japan |
| *P. schewiakoffi* | G4_Psche | NA |
| *P. decaurelia* | decaurelia_223 | NA |
| *P. dodecaurelia* | dodecaurelia_274 | NA |
| *P. jenningsi* | jenningsi_M | NA |
| *P. novaurelia* | novaurelia_TE | NA |
| *P. octaurelia* | octaurelia_138 | NA |
| *P. pentaurelia* | pentaurelia_87 | NA |
| *P. sonneborni* | sonneborni_30995 | NA |
| *P. calkinsi* | CL13-6_calkinski | NA |
| *P. octaurelia* | Brazil_P11_octaurelia | Brazil |

**Fig. S1.**
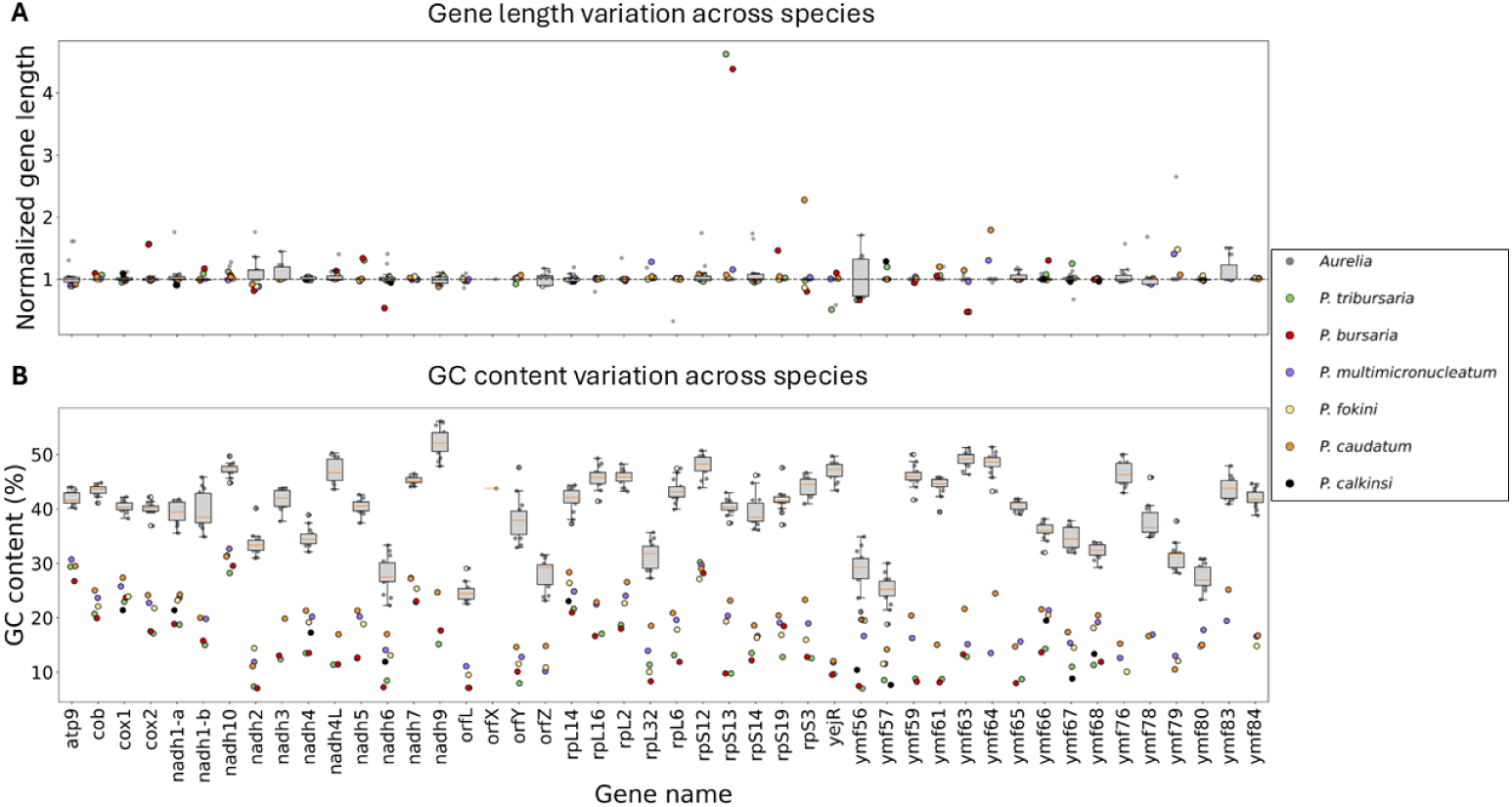
(A) Normalized gene length across mitochondrial genes. (B) GC content (%) across genes. Gray points indicate *Paramecium aurelia* species; colored points represent outgroups.

**Table S2.** Majority-vote gene-tree support for key backbone relationships in *Paramecium*.

| Gene | Bursaria<br>outgroup | Multi-Foki<br>clade together | Caudatum<br>outgroup | Sexaurelia<br>outgroup | Aurelia<br>clade together |
| --- | --- | --- | --- | --- | --- |
| atp9 | Yes | Yes | No | No | No |
| cob | Yes | Yes | Yes | Yes | Yes |
| cox1 | Yes | Yes | Yes | Yes | Yes |
| cox2 | Yes | Yes | Yes | No | Yes |
| nadh10 | Yes | Yes | No | Yes | Yes |
| nadh1-a | Yes | Yes | Yes | Yes | Yes |
| nadh1-b | Yes | Yes | Yes | No | Yes |
| nadh2 | Yes | Yes | Yes | Yes | Yes |
| nadh3 | Yes | NA | Yes | No | Yes |
| nadh4 | Yes | Yes | Yes | Yes | Yes |
| nadh4L | Yes | Yes | Yes | No | Yes |
| nadh5 | Yes | Yes | Yes | Yes | Yes |
| nadh6 | Yes | Yes | Yes | No | Yes |
| nadh7 | Yes | Yes | Yes | No | Yes |
| nadh9 | Yes | NA | Yes | No | Yes |
| rpL14 | Yes | Yes | Yes | Yes | Yes |
| rpL16 | Yes | Yes | Yes | No | Yes |
| rpL2 | Yes | Yes | Yes | No | Yes |
| rpL32 | Yes | Yes | Yes | No | Yes |
| rpL6 | Yes | Yes | Yes | Yes | Yes |
| rpS12 | Yes | Yes | Yes | Yes | Yes |
| rpS13 | Yes | Yes | Yes | No | Yes |
| rpS14 | Yes | Yes | Yes | Yes | Yes |
| rpS19 | Yes | Yes | Yes | Yes | Yes |
| rpS3 | Yes | Yes | Yes | Yes | Yes |
| yejR | Yes | Yes | Yes | Yes | Yes |
| ymf56 | Yes | Yes | Yes | No | Yes |
| ymf57 | Yes | Yes | Yes | Yes | Yes |
| ymf59 | Yes | Yes | Yes | Yes | Yes |
| ymf61 | Yes | Yes | Yes | No | Yes |
| ymf63 | Yes | Yes | Yes | Yes | Yes |
| ymf64 | NA | Yes | Yes | No | Yes |
| ymf65 | Yes | Yes | Yes | Yes | Yes |
| ymf66 | Yes | Yes | Yes | Yes | Yes |
| ymf67 | Yes | Yes | Yes | Yes | Yes |
| ymf68 | Yes | Yes | Yes | Yes | Yes |
| ymf76 | Yes | Yes | Yes | Yes | Yes |
| ymf78 | NA | Yes | Yes | No | Yes |
| ymf79 | NA | Yes | Yes | No | Yes |
| ymf80 | NA | Yes | Yes | Yes | Yes |
| ymf83 | NA | Yes | Yes | Yes | Yes |
| ymf84 | NA | Yes | Yes | No | Yes |

**Fig. S2.**
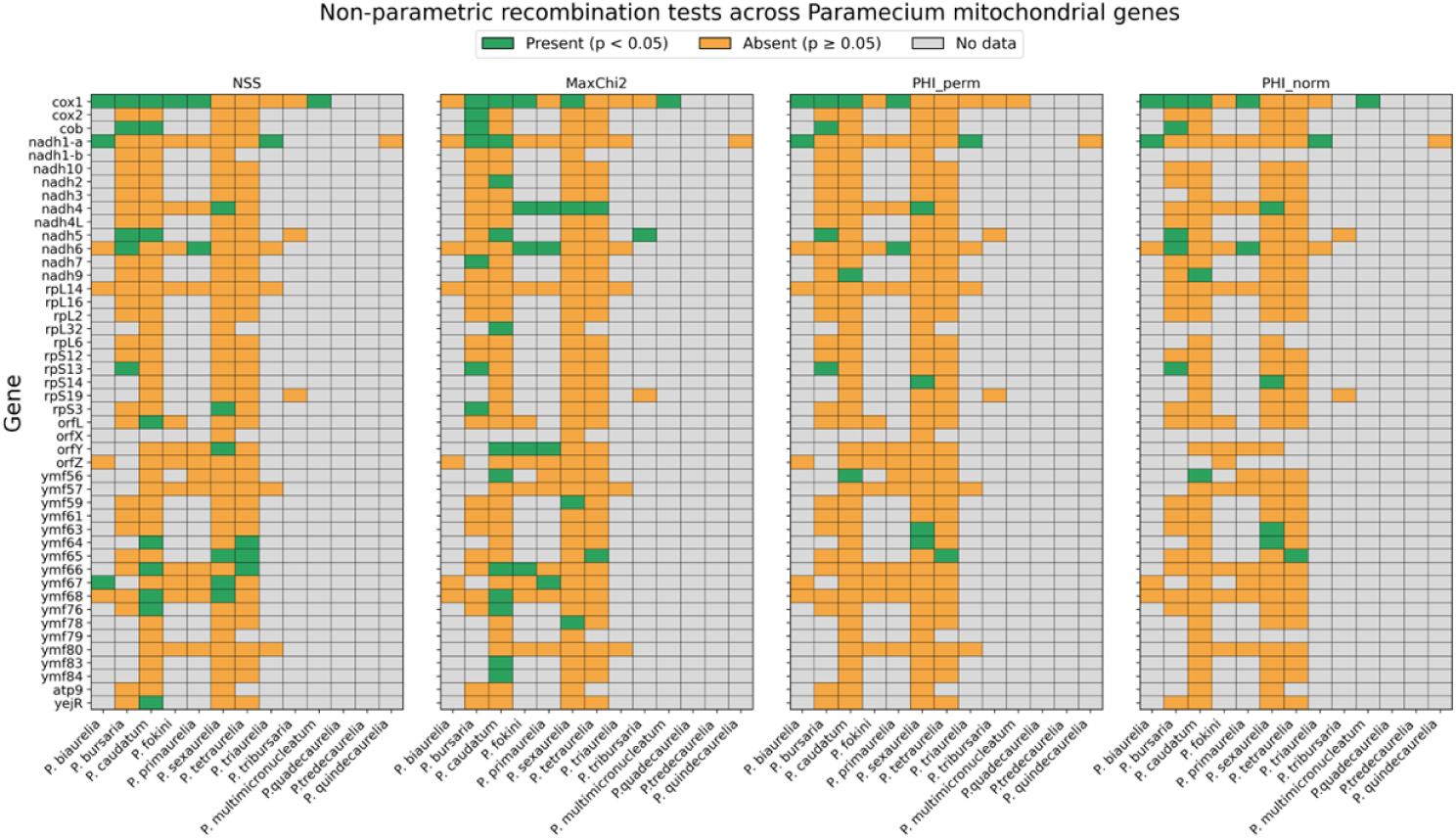
Genome-wide heatmaps of non-parametric recombination tests across *Paramecium* mitochondrial genes. NSS, MaxChi2, PHI perm, and PHI norm are shown side-by-side. Green indicates significant recombination (p ¡ 0.05), orange indicates no evidence (p *≥* 0.05), and gray denotes missing data.

**Fig. S3.**
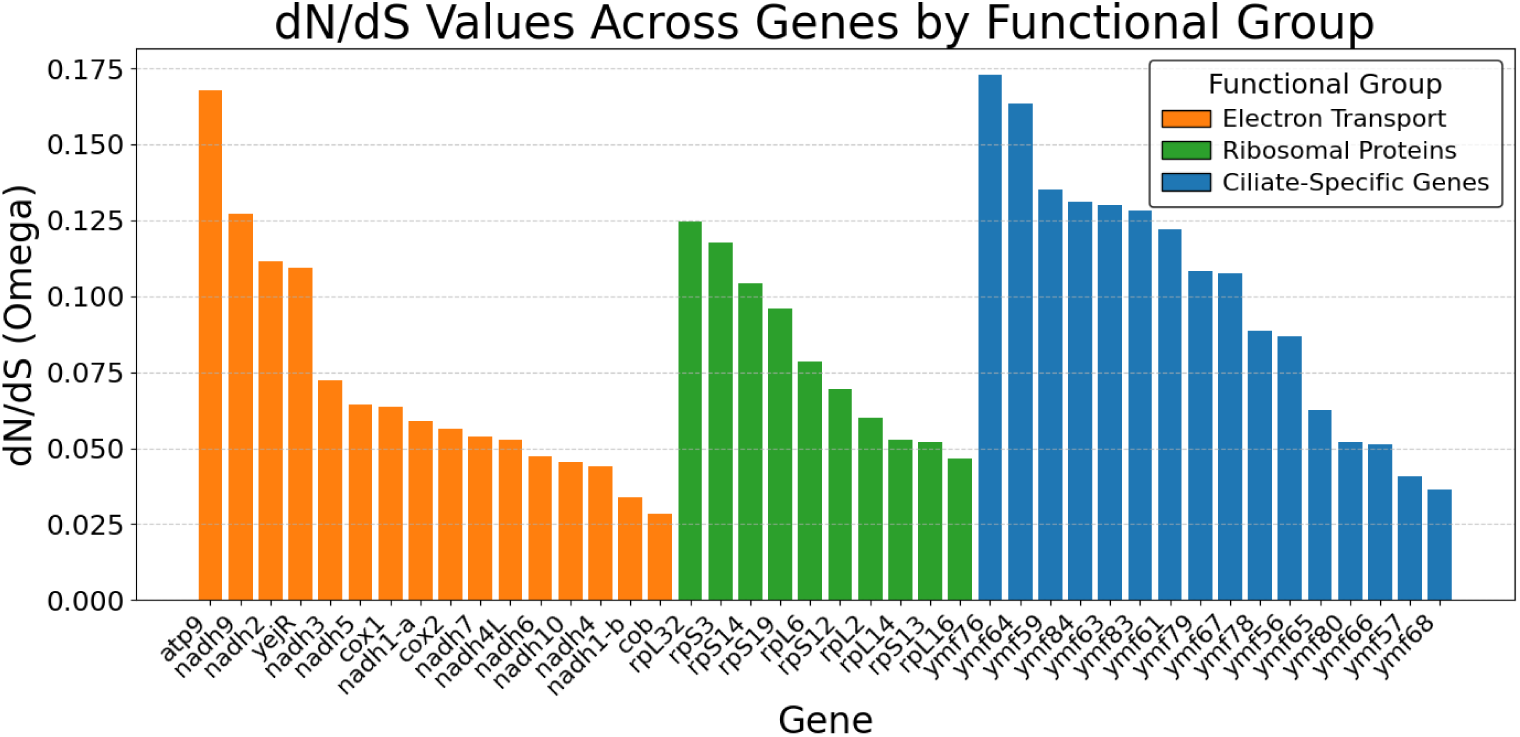
Gene-wise dN/dS (*ω*) estimates across mitochondrial protein-coding genes in *Paramecium*. Genes are grouped by functional category: electron transport chain (ETC), ribosomal proteins (LSU/SSU), and ciliate-specific (ymf) genes. Each bar represents the dN/dS value for an individual gene.

**Table S3.**
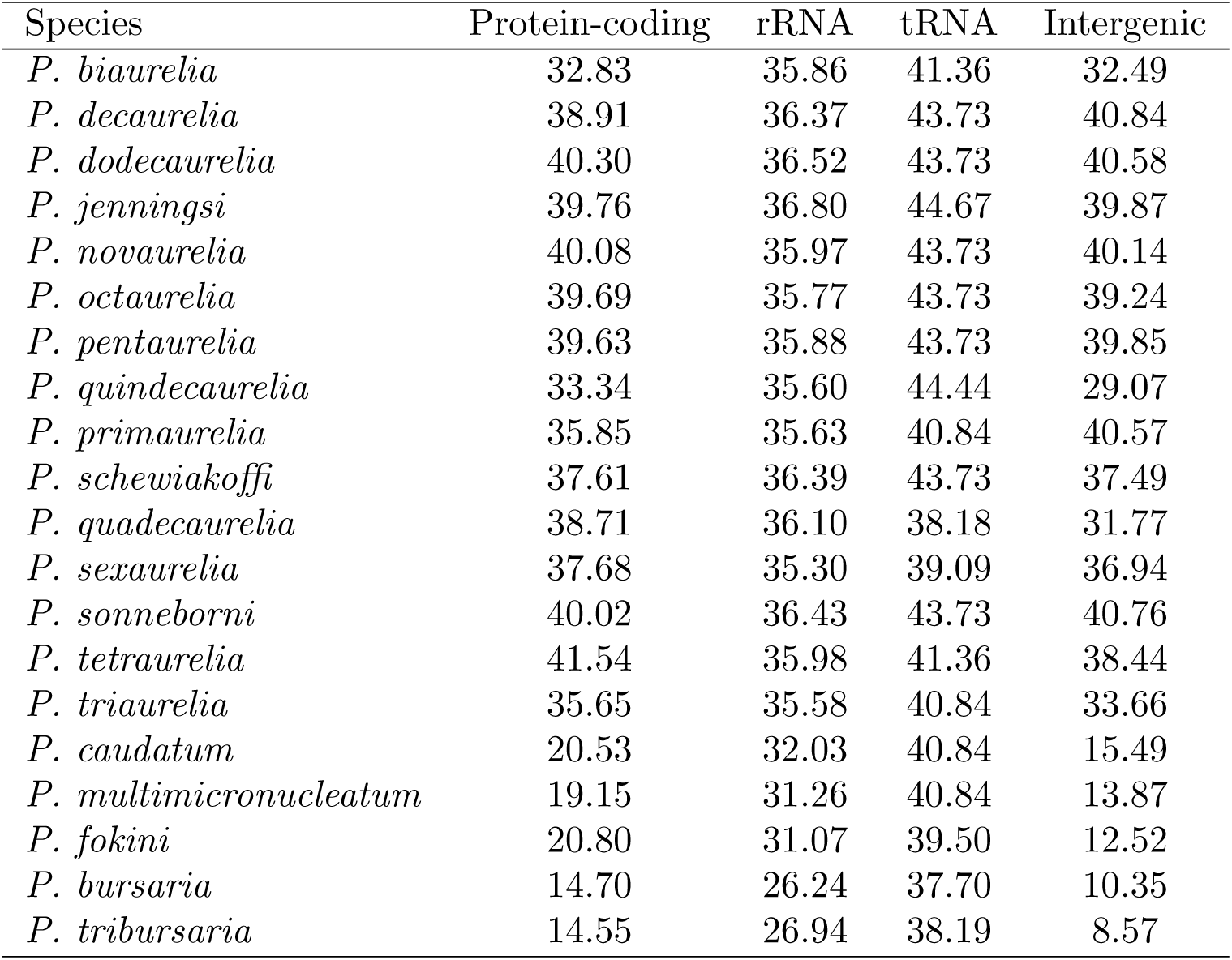
GC content (%) of protein-coding genes, rRNAs, tRNAs, and intergenic regions across *Paramecium* mitochondrial genomes.

**Table S4.** Evidence for recombination based on gene conversion across *Paramecium* species. Estimates are shown for different values of *θ*.

| Species | $\theta$ | Composite likelihood | G4 | $R^2$ distance | $D'$ distance | $4N_e r$ | $4N_e r$ |
| --- | --- | --- | --- | --- | --- | --- | --- |
|  |  | p-value | p-value | p-value | p-value | (Hudson) | (Wakeley) |
| *P. biaurelia* | 0.001 | 0.000 | 0.125 | 0.000 | 0.113 | 3 | 1.769 |
|  | 0.01 | 0.000 | 0.114 | 0.000 | 0.097 | 3 | 1.769 |
| *P. bursaria* | 0.01 | 0.193 | 0.971 | 0.479 | 0.943 | 0 | 0 |
|  | 0.1 | 0.590 | 0.966 | 0.507 | 0.919 | 0 | 0 |
| *P. caudatum* | 0.01 | 0.739 | 0.948 | 0.024 | 0.742 | 0 | 0 |
|  | 0.1 | 0.880 | 0.951 | 0.026 | 0.778 | 0 | 0 |
| *P. multimicronucleatum* | 0.001 | 0.853 | 0.223 | 0.325 | 0.217 | 1 | 0 |
|  | 0.01 | 0.629 | 0.218 | 0.295 | 0.210 | 0 | 0 |
| *P. fokini* | 0.01 | 0.694 | 1.000 | 0.852 | 1.000 | 0 | 0 |
|  | 0.1 | 0.015 | 1.000 | 0.843 | 1.000 | 0 | 0 |
| *P. quindecarelia* | 0.01 | 0.784 | 1.000 | 0.001 | 1.000 | 0 | 0 |
|  | 0.1 | 0.944 | 1.000 | 0.003 | 1.000 | 0 | 0 |
| *P. primaurelia* | 0.01 | 0.982 | 0.165 | 0.000 | 0.009 | 1 | 0 |
|  | 0.1 | 0.829 | 0.158 | 0.000 | 0.009 | 1 | 0 |
| *P. sexaurelia* | 0.01 | 1.000 | 0.700 | 0.286 | 0.678 | 0 | 2.173 |
|  | 0.1 | 0.990 | 0.725 | 0.310 | 0.708 | 0 | 2.173 |
| *P. tetraurelia* | 0.001 | 0.302 | 0.587 | 0.866 | 0.565 | 0 | 0 |
|  | 0.01 | 0.400 | 0.596 | 0.849 | 0.568 | 0 | 0 |
| *P. triaurelia* | 0.001 | 0.805 | 0.786 | 0.076 | 0.597 | 0 | 0 |
|  | 0.01 | 0.555 | 0.809 | 0.074 | 0.623 | 0 | 0 |
| *P. tribursaria* | 0.001 | 1.000 | 0.979 | 0.996 | 0.979 | 0 | 0 |
|  | 0.01 | 0.869 | 0.979 | 0.999 | 0.979 | 0 | 0 |

**Table S5.**
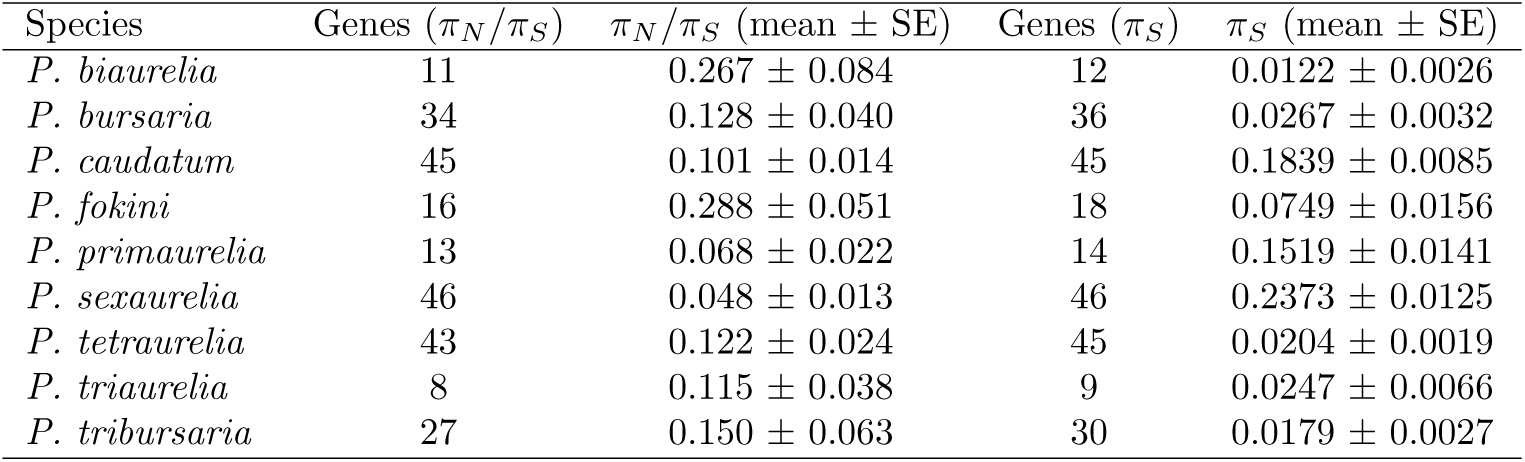
Per-species estimates of nonsynonymous to synonymous diversity (*π_N_ /π_S_*) and synonymous diversity (*π_S_*) across *Paramecium* mitochondrial genes. Values are means across genes *±* standard error (SE). Species with fewer than five genes were excluded from this analysis.

| Species | Genes ($\pi_N/\pi_S$) | $\pi_N/\pi_S$ (mean ± SE) | Genes ($\pi_S$) | $\pi_S$ (mean ± SE) |
| --- | --- | --- | --- | --- |
| *P. biaurelia* | 11 | 0.267 ± 0.084 | 12 | 0.0122 ± 0.0026 |
| *P. bursaria* | 34 | 0.128 ± 0.040 | 36 | 0.0267 ± 0.0032 |
| *P. caudatum* | 45 | 0.101 ± 0.014 | 45 | 0.1839 ± 0.0085 |
| *P. fokini* | 16 | 0.288 ± 0.051 | 18 | 0.0749 ± 0.0156 |
| *P. primaurelia* | 13 | 0.068 ± 0.022 | 14 | 0.1519 ± 0.0141 |
| *P. sexaurelia* | 46 | 0.048 ± 0.013 | 46 | 0.2373 ± 0.0125 |
| *P. tetraurelia* | 43 | 0.122 ± 0.024 | 45 | 0.0204 ± 0.0019 |
| *P. triaurelia* | 8 | 0.115 ± 0.038 | 9 | 0.0247 ± 0.0066 |
| *P. tribursaria* | 27 | 0.150 ± 0.063 | 30 | 0.0179 ± 0.0027 |

**Table S6.**
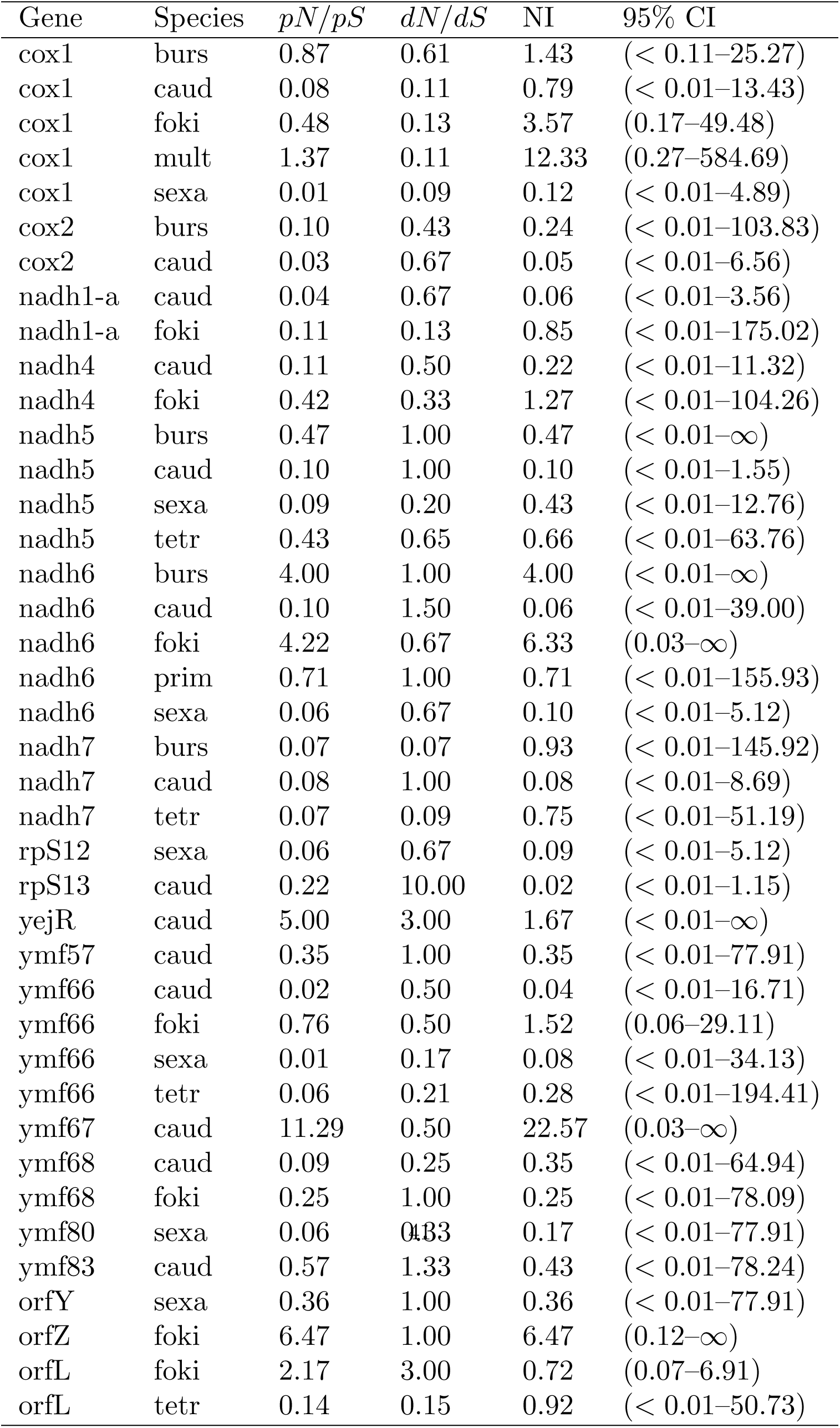
Neutrality index (NI) estimates across mitochondrial genes. Values are shown for gene–species pairs, including polymorphism (*pN/pS*), divergence (*dN/dS*), neutrality index (NI), and 95% confidence intervals. Wide confidence intervals reflect limited polymorphism and small denominators in some comparisons.

**Table S7.**
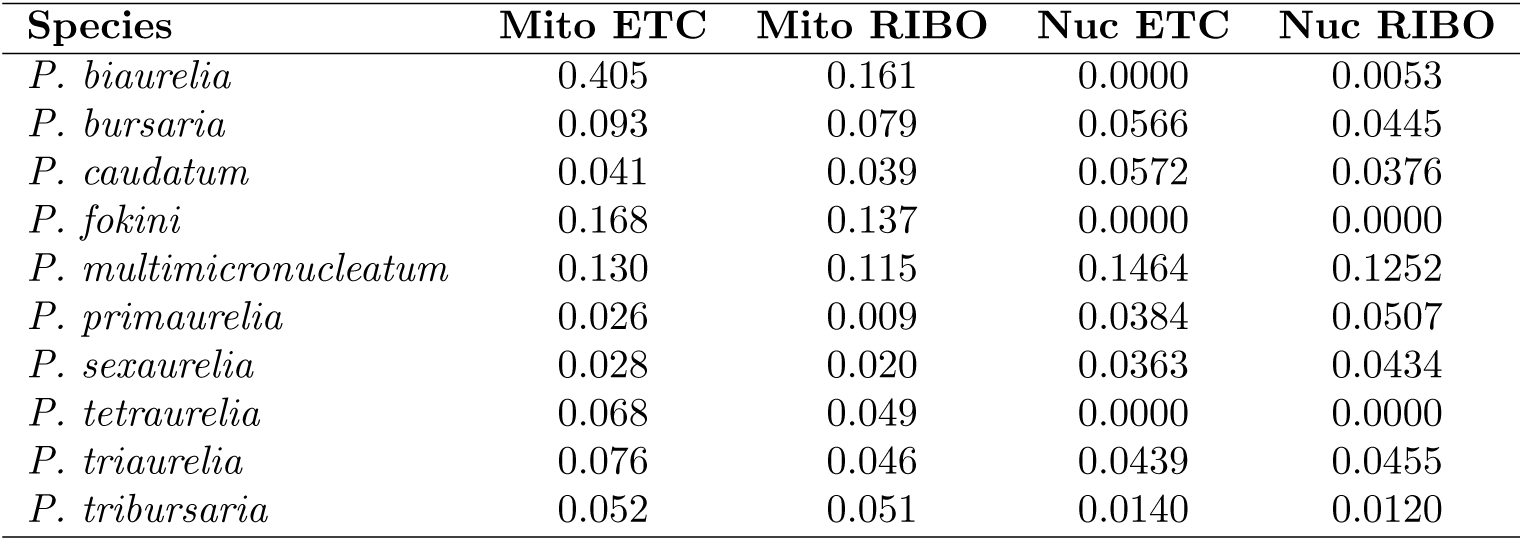
Median *π_N_ /π_S_* across species for electron transport chain (ETC) and ribosomal (RIBO) genes in mitochondrial and nuclear genomes.

**Table S8.** Relative effective population size 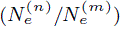 and mutation-rate ratios 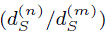 across *Paramecium* species.

| Species | $N_e^{(n)}/N_e^{(m)}$ | $d_S^{(n)}/d_S^{(m)}$ |
| --- | --- | --- |
| *P. biaurelia* | 1.03 | 0.31 |
| *P. primaurelia* | 0.90 | 0.45 |
| *P. sexaurelia* | 0.10 | 0.49 |
| *P. tetraurelia* | 0.53 | 0.31 |

**Table S9.**
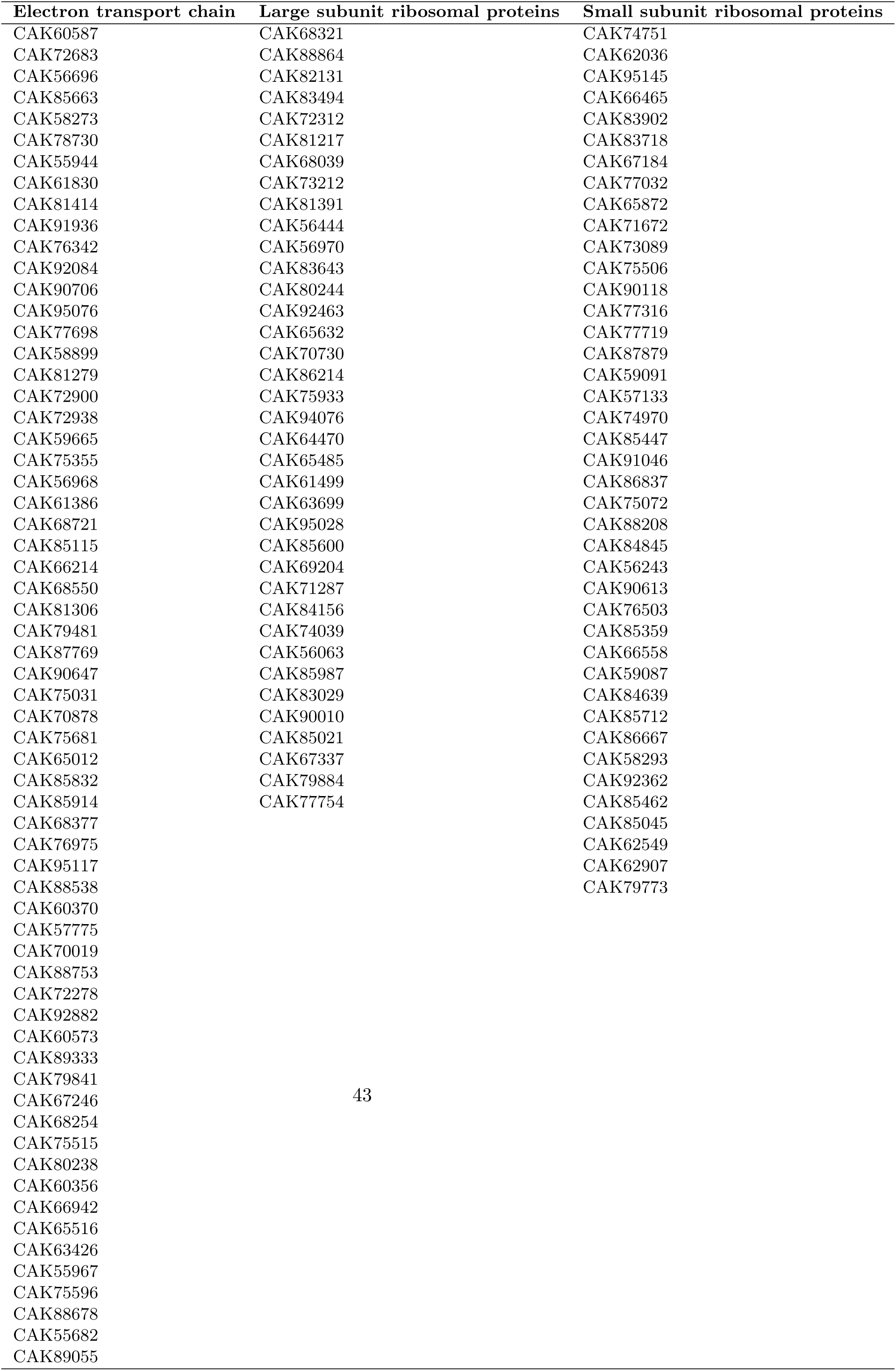
Nuclear-encoded gene IDs for functional groups.

## Note 1. Unannotated ORFs in *P. bursaria* mitochondrial genomes

To examine the origin of apparent gaps in *P. bursaria* mitochondrial gene organization, we identified unannotated ORFs ≥ 100 aa after excluding annotated genes and overlapping regions, retaining the longest ORF at each locus. These ORFs were classified into conserved (“R”) and lineage-restricted (“G”) clusters using all-versus-all BLASTP (≥ 30% identity, ≥ 60% coverage). Conserved ORFs occupy many of the apparent gaps, indicating that these regions correspond to previously unrecognized genes rather than nonfunctional intergenic sequence.

The mitochondrial genomes of *P. bursaria* harbor numerous recurring lineage-restricted ORFs (e.g., G6, G8, G13) that are consistently detected across isolates, indicating that they represent authentic genomic elements rather than artifacts of annotation. Although some of these ORFs show weak similarity to *ymf*-like genes, these matches are inconsistent, suggesting that they represent highly diverged homologs that are difficult to detect using standard sequence similarity approaches. Consistent with this, these G clusters exhibit elevated within-species polymorphism relative to core mitochondrial genes, with values of G1 ≈ 0.12, G22 ≈ 0.18, G25 ≈ 0.39, G13 ≈ 0.50, G8 ≈ 0.63, G14 ≈ 0.63, and G6 ≈ 1.03 (mean ≈ 0.43), consistent with relaxed functional constraint.

## Notes

### Competing Interest Statement

The authors have declared no competing interest.

